# The MRAP2 accessory protein directly interacts with melanocortin-3 receptor to enhance signaling

**DOI:** 10.1101/2024.11.06.622243

**Authors:** Aqfan Jamaluddin, Rachael A. Wyatt, Joon Lee, Georgina K.C. Dowsett, John A. Tadross, Johannes Broichhagen, Giles S.H. Yeo, Joshua Levitz, Caroline M. Gorvin

## Abstract

The central melanocortin system links nutrition to energy expenditure, with melanocortin-4 receptor (MC4R) controlling appetite and food intake, and MC3R regulating timing of sexual maturation, rate of linear growth and lean mass accumulation. Melanocortin-2 receptor accessory protein-2 (MRAP2) is a single transmembrane protein that interacts with MC4R to potentiate it’s signalling, and human mutations in MRAP2 cause obesity. Previous studies have been unable to consistently show whether MRAP2 affects MC3R activity. Here we used single-molecule pull-down (SiMPull) to confirm that MC3R and MRAP2 interact in HEK293 cells. Analysis of fluorescent photobleaching steps showed that MC3R and MRAP2 readily form heterodimers most commonly with a 1:1 stoichiometry. Human single-nucleus and spatial transcriptomics show MRAP2 is co-expressed with MC3R in hypothalamic neurons with important roles in energy homeostasis and appetite control. Functional analyses showed MRAP2 enhances MC3R cAMP signalling, impairs β-arrestin recruitment, and reduces internalization in HEK293 cells. Structural homology models revealed putative interactions between the two proteins and alanine mutagenesis of five MRAP2 and three MC3R transmembrane residues significantly reduced MRAP2 effects on MC3R signalling. Finally, we showed genetic variants in MRAP2 that have been identified in individuals that are overweight or obese prevent MRAP2’s enhancement of MC3R-driven signalling. Thus, these studies reveal MRAP2 as an important regulator of MC3R function and provide further evidence for the crucial role of MRAP2 in energy homeostasis.

## Introduction

The melanocortin receptor-2 accessory protein 2 (MRAP2) is a single-pass transmembrane protein that modulates the function of several G protein-coupled receptors (GPCRs) expressed in the hypothalamus that regulate food intake (*1–4*). These GPCRs include melanocortin receptor-4 (MC4R), a central regulator of appetite, inactivating mutations of which are the most common genetic cause of obesity, and the receptor for ghrelin (growth hormone secretagogue receptor, GHSR), which enhances appetite (*1, 2, 4*). Similarly to MC4R, human genetic variants in MRAP2 have been identified in several families and individuals with obesity and reduce MC4R activity (*5–7*). MRAP2 was identified as a homolog of MRAP1, an accessory protein that is essential for the cell-surface expression and ligand responsiveness of melanocortin receptor-2 (MC2R), which regulates adrenal development and steroidogenesis (*2*). Unlike MRAP1, the MRAP2 protein is not essential for GPCR expression at the cell surface. However, MRAP2 enhances MC4R expression at neuronal primary cilia, a microtubule-based organelle with a vital role in appetite regulation (*8*), suggesting that MRAP2 may establish signaling hubs that favour receptor signaling.

Deletion of the MRAP2 gene from mice on a variety of genetic backgrounds is associated with extreme obesity, increased fat mass and visceral adiposity, analogous to MC4R knockout mice (*9, 10*). Double knockouts of MRAP2 and MC4R demonstrate that MRAP2 facilitates the action of MC4R, but that there are also MC4R-independent mechanisms (*5*). MRAP2 mice lack the early-onset hyperphagia of MC4R knockout mice, and humans with MRAP2 genetic variants exhibit hyperglycaemia, hypertension and high blood cholesterol more frequently than those with MC4R mutations (*6*). This is consistent with studies showing that MRAP2 can modulate the signaling profile of several GPCRs involved in energy homeostasis. Thus, MRAP2 enhances signaling by MC4R and the ghrelin receptor, while it suppresses the activity of the prokineticin receptors (*3*), orexin receptor-1 (*11*) and melanin concentrating hormone receptor-1 (*12*). One study identified >40 putative binding partners for MRAP2 (*13*); however, signaling data was not provided for most receptors, and some had previously been described as non-interacting proteins, therefore further work is required to validate these findings. Additionally, while several studies have shown that MC4R signaling is impaired by some MRAP2 genetic variants identified in overweight or obese individuals (*6, 7, 14*), their effect on signaling by other MRAP2 interacting proteins remains to be explored.

Co-immunoprecipitation studies have shown that MRAP2 can interact with all five members of the melanocortin receptor family when overexpressed in cell-lines (*2, 15*). MC3R is a negative regulator of the central melanocortin system (*16, 17*). It is required for the normal activation of AgRP neurons in response to nutritional deficit (*16*). Deletion of MRAP2 from AgRP neurons also blunts their fasting-induced activation (*1*), similarly to MC3R, and it has been hypothesized that a complex signaling system may exist between MC3R, MRAP2 and other receptors at these neurons (*16*). There is some evidence that MC3R may interact with MRAP2, although this is inconclusive. MRAP2 coimmunoprecipitates with MC3R (*2*) and enhances ciliary expression of the receptor in transfected cells (*8*). However, co-expression of MC3R and MRAP2 has been shown to reduce cAMP signaling (*2*), enhance signaling (*5*), or have no effect on signaling (*18, 19*). This motivates a more comprehensive examination of the effect of MRAP2 on MC3R activity. Such inconsistencies are common in the MRAP2 literature, including for MC4R, with MRAP2 initially described to reduce MC4R cell surface expression and impair it’s signaling, then later shown to increase MC4R function, consistent with mouse knockout studies (*2, 3, 13*). These discrepancies are likely due to large variations in studies seeking to investigate MRAP2 function. These include overexpressing MRAP2 at DNA ratios of 3-20x that of GPCR, although no rationale is provided for these experimental decisions (*11–13, 20*). As such, these high concentrations of MRAP2 could lead to overexpression artefacts and false positive results (*21*). A recent preprint demonstrated that MRAP2 is still capable of enhancing MC4R signaling when the two proteins are expressed at equal concentrations, and that MRAP2 overexpression may affect GPCR oligomer assembly (*22*), indicating that studies of equal concentrations of MRAP2 and GPCRs are required to ensure that molecular details are not missed.

MRAP2 facilitates signaling by some GPCRs (*5, 20*) and suppresses responses by other receptors (*3, 22*). Studies focussed predominantly on the ghrelin receptor have elucidated several mechanisms by which MRAP2 may enhance signaling. These include a reduced ability to recruit β-arrestin proteins albeit with no change in receptor cell surface expression (*20*). A similar mechanism has been suggested for the Prokineticin Receptor-2 (*23*) and MC4R (*20*). Additionally, MRAP2 biases GHSR signaling to reduce Rho activation, enhances G protein coupling of MC4R, and may reduce MC4R oligomerization that can suppress receptor signaling (*20, 22*). The structural regions involved in MRAP2 interaction with GPCRs remain largely unexplored. MC4R homology models based on the cryo-EM structure of the MC2R-MRAP1 complex suggest that MRAP2 may interact with transmembrane helix (TM)-5 or TM6, but no mechanistic studies were performed (*20*). Additionally, while large truncation mutations (e.g. deletion of the transmembrane region, deletion of the C-tail) of MRAP2 show loss of interaction or impaired signaling (*11*), these do not provide insights into the specific residues involved or their mechanisms of action.

Here we examined the effect of MRAP2 on MC3R activity in HEK293 cells. We demonstrated that MRAP2 interacts with MC3R in a 1:1 dimer to enhance cAMP signaling, reduce β-arrestin recruitment and impair receptor internalization. Structural homology models and alanine mutagenesis identified critical residues important for the interaction. Finally, we demonstrated that MRAP2 variants identified in individuals that are overweight or obese reduce MC3R signaling and enhance receptor internalization.

## Results

### MRAP2 is colocalised with MC3R in neurons involved in energy homeostasis

Previous studies have been unable to determine whether MRAP2 interacts with MC3R to influence receptor signaling (*2, 5, 18, 19*). As co-expression in the same cells is a requirement for biologically relevant MC3R-MRAP2 interactions, we first assessed expression of the transcripts encoding MC3R and MRAP2 proteins in HYPOMAP, a single-nucleus RNA sequencing (snRNAseq) and spatial transcriptomic atlas of the human hypothalamus (*24*). snRNA-seq data allowed quantification of the expression of the two genes in neuronal cells. MRAP2 was expressed in ∼35% of all neuronal cells and was detected in 57% of MC3R-positive neurons indicating that the two proteins have some co-expression in physiologically relevant cell types (Figure 1, Table S1). By comparison, in HYPOMAP, MRAP2 was detected in 53% of MC4R-positive neurons (Figure 1, Table S1). Visium spatial transcriptomics revealed that MRAP2 is expressed throughout the hypothalamus, particularly in regions where there is greater neuronal density, whereas MC3R expression is more restricted to the arcuate nucleus, ventromedial hypothalamus and periventricular region (Figure S1, Table S1). MRAP2 transcripts are present under the same spatially barcoded spots as *MC3R* transcripts in these regions, which are known to have important roles in energy homeostasis and appetite control.

**Figure 1.**
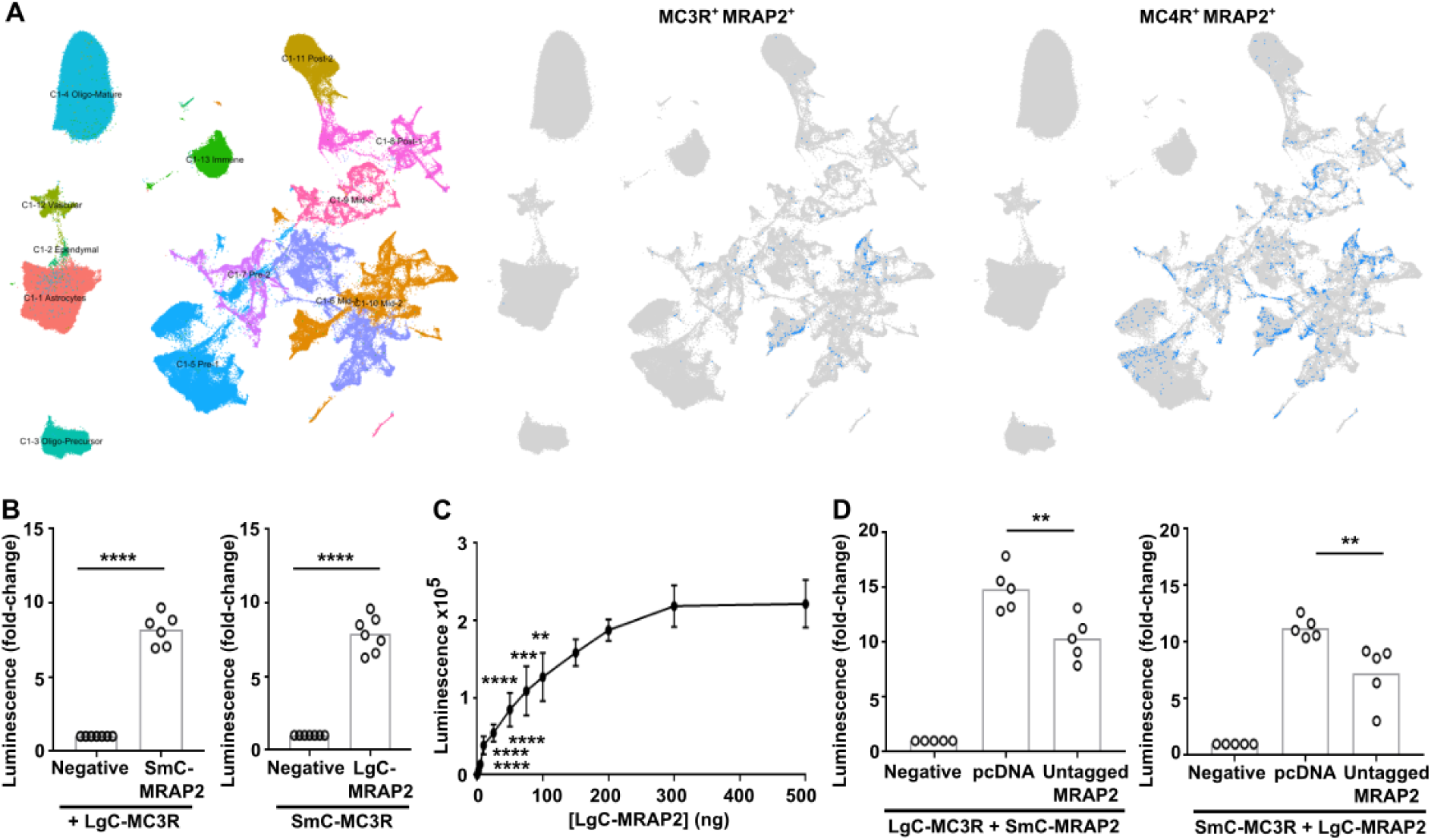
MC3R is co-expressed with MRAP2 in hypothalamic neurons. (**A**) snRNAseq of the human hypothalamus reveals co-expression of MRAP2 with MC3R and MC4R. (Left) UMAP plot of the snRNAseq data from HYPOMAP, with cells coloured by C1 clustering. (Middle) UMAP plot highlighting cells in blue that co-express MRAP2 and MC3R. (Right) UMAP plot highlighting cells in blue that co-express MRAP2 and MC4R transcripts. Table S1 shows the top 15 clusters with the highest MC3R expression or MC4R expression, with the percentage co-expression of MRAP2 in each cluster. (**B**) NanoBiT luminescence between MC3R and MRAP2 or negative control. N=6-7. (**C**) NanoBiT luminescence between 100ng SmC-MC3R and increasing concentrations of LgC-MRAP2. N=4. (**D**) Competition assays with NanoBiT constructs and pcDNA or MRAP2. N=5. Statistical analyses performed with student’s t-test in B, one-way ANOVA with Dunnett’s test in D. ****p<0.0001, ***p<0.001, **p<0.01.

### MRAP2 interacts with MC3R

Our studies have shown that MC3R colocalises with MRAP2 in hypothalamic neurons that are known to regulate energy homeostasis. To determine whether MC3R and MRAP2 are likely to interact we first assessed protein proximity in transiently transfected HEK293 cells using the NanoBiT split-luciferase system with both proteins tagged at the C-terminus. There was increased luminescence observed in cells co-expressing MC3R and MRAP2 (100 ng each) compared to cells expressing MC3R and the negative control (Figure 1B). Similar luminescence values were observed in cells transfected with either iteration of NanoBiT tags (i.e. LgC-MC3R and SmC-MRAP2 or SmC-MC3R and LgC-MRAP2). Saturation curves were performed in which a fixed amount of MC3R (100 ng) was transfected with increasing concentrations of MRAP2. This showed a hyperbolic increase in the luminescence indicating the signal is unlikely to be due to random collisions (Figure 1C). Co-transfection of cells with untagged MRAP2 to compete with SmC/LgC-MRAP2 reduced NanoBiT luminescence values (Figure 1D), providing further evidence that the two proteins may interact.

Although NanoBiT assays can indicate proximity between proteins, these assays do not measure interactions with single complex precision and cannot accurately measure stoichiometry. To assess this and verify the interaction, we used the single-molecule pull-down (SiMPull) technique, which has previously been used to assess heteromeric GPCR complexes (*25*) (Figure 2A). We generated MC3R and MRAP2 constructs with N-terminal hemagglutinin (HA) or FLAG epitopes followed by a SNAP, Halo or CLIP tag amenable to labelling with organic dyes (Figure S2-S3, Table S2), and demonstrated that MC3R maintained receptor function, MC3R and MRAP2 colocalized in cells when transiently transfected and MRAP2 could enhance signaling by a known interacting receptor, MC4R (Figure S2-S3). We first used SiMPull to determine the expression and stoichiometry of MC3R homomers. Cells were transfected with HA-Halo-MC3R and labelled with membrane impermeable CA-Sulfo646 (*26*), then cells were lysed, receptors immobilized by anti-HA antibodies and single molecules imaged by total internal reflection fluorescence microscopy. The majority of molecules showed single bleaching steps (∼83%), while approximately 15% had two steps per molecule (Figure 2B-D), indicating that most MC3R is monomeric at the cell surface. In the absence of anti-HA antibodies very few molecules (6 molecules across 5 images) were observed (Figure S4). We also examined MRAP2 stoichiometry by SiMPull as it has been described to form homodimers or higher-order oligomers in several studies (*2, 27, 28*). We first verified that the known dimeric GPCR mGluR2 produced single-molecules with two photobleaching steps (*29*) (Figure S4). Cells were then transfected with HA-Halo-MRAP2, labelled with CA-Sulfo646 and imaged. MRAP2 showed single bleaching steps in ∼68% of molecules, while ∼28% had two bleaching steps, indicating some dimer formation may occur. A small number of molecules (<5%) had three or four bleaching steps corresponding to higher-order oligomers (Figure 2E-G). Therefore, MRAP2 primarily forms stable monomers or dimers when expressed alone. To assess MC3R and MRAP2 heteromers, HA-Halo-MC3R and FLAG-CLIP-MRAP2 were transfected in HEK293 cells and Halo and CLIP tags labelled with CA-Sulfo646 and BC-DY547 fluorophores, respectively, prior to lysis. Receptors were immobilized by anti-HA antibodies and fluorescence co-localization was assessed. In the absence of MC3R, there were negligible single molecules observed (13 molecules across 5 images) (Figure 2H). In co-transfected cells, co-localization was present in almost 30% of MC3R spots (Figure S4). Photobleaching step analysis showed 1-step each for MC3R and MRAP2 in ∼74% of co-localized spots, while some 2- and 3-step bleaching was observed for MRAP2 (Figure 2I-J). Less than 5% of spots showed two MC3R and two MRAP2 bleaching steps. To verify these findings the SiMPull experiments were repeated with the Halo and CLIP labels swapped. Thus, cells were transfected with HA-Halo-MRAP2 and FLAG-CLIP-MC3R, then labeled and imaged as described. FLAG-CLIP-MC3R expression alone produced few single molecules (24 molecules across 5 images) (Figure S4). These studies had a similar total number of co-localized spots (∼32% of receptor spots). Bleaching step analysis of these spots showed 63% had one MC3R and one MRAP2 step, while 21% had two MRAP2 steps, ∼7% had 3 steps for MRAP2, and ∼7.5% had two steps each for MC3R and MRAP2 (Figure S4). In contrast, MRAP2 did not pull-down or colocalize with SSTR3, a receptor that is not known to interact with MRAP2 and whose signaling is not enhanced by MRAP2 (Figure S5). These studies indicate that MC3R is more likely to interact with MRAP2 in a 1:1 stoichiometry but can interact with more than one MRAP2 molecule.

**Figure 2.**
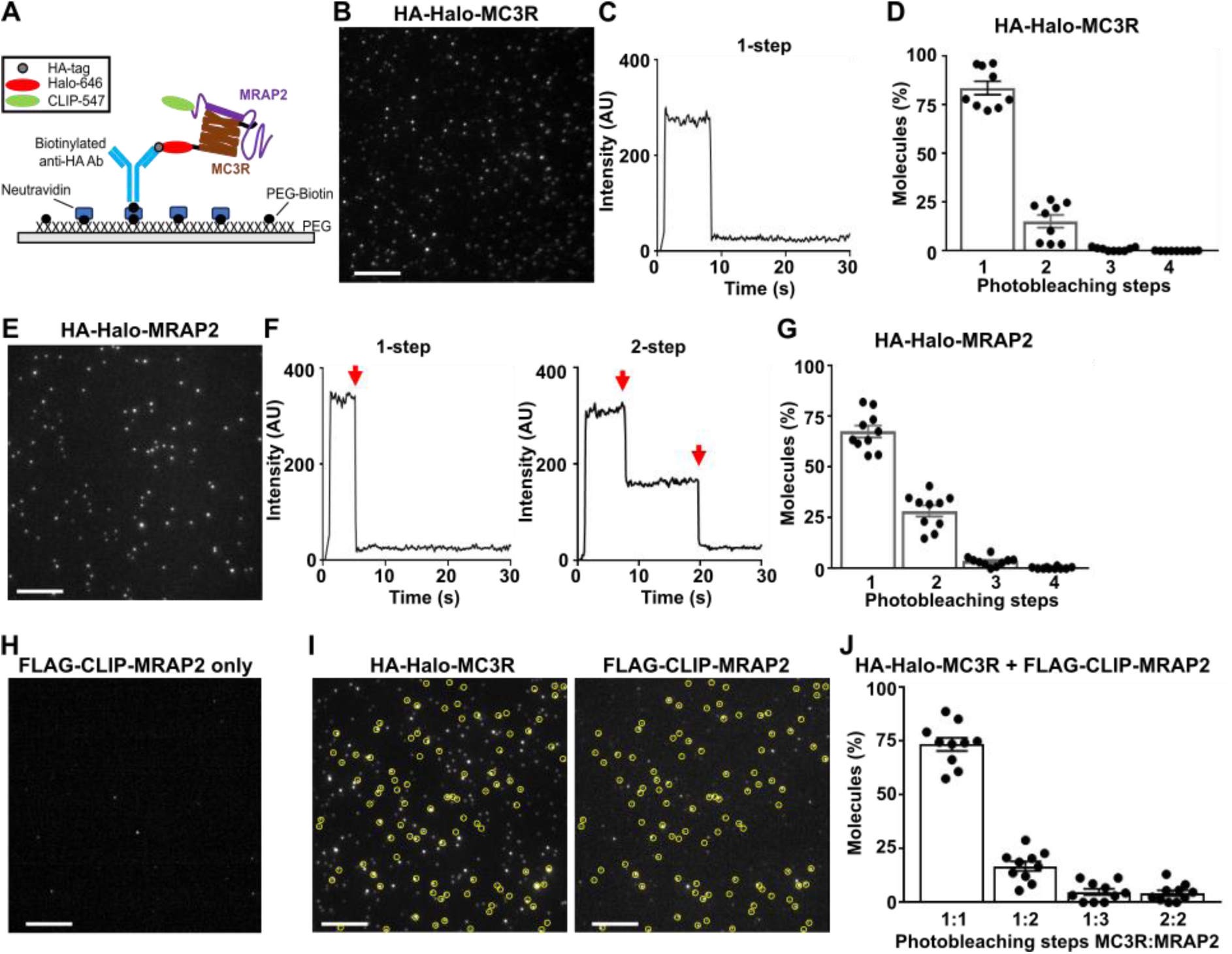
MC3R and MRAP2 interact primarily in a 1:1 stoichiometry. (**A**) Schematic of two-color SiMPull experiments. Fresh cell lysate from HEK293 cells expressing HA-Halo-MC3R with FLAG-CLIP-MRAP2 is added to a PEG-passivated glass slide containing immobilized anti-HA antibody. Halo and CLIP tags are labeled with CA-Sulfo646 and BC-DY547, respectively. (**B**) Representative single-molecule fluorescence image of HA-Halo-MC3R with (**C**) examples of single-molecule fluorescence traces with photobleaching steps (red arrows). (**D**) Proportion of molecules with 1 to 4 bleaching steps. N = 1565 molecules from 10 movies. (**E**) Representative single-molecule fluorescence image of HA-Halo-MRAP2 with (**F**) examples of single-molecule fluorescence traces with photobleaching steps (red arrows). (**G**) Quantification of molecules with 1 to 4 bleaching steps. N = 1333 molecules from 10 movies. (**H**) Cells transfected with FLAG-CLIP-MRAP2 only, showing negligible background fluorescence. (**I**) Representative two-color SiMPull images of HA-Halo-MC3R and FLAG-CLIP-MRAP2 with colocalized spots circled in yellow, and (**J**) Photobleaching step analysis from colocalized spots showing MC3R interacts with MRAP2 monomers, and occasionally dimers. N=926 molecules from 10 movies. Scale, 10 μm for all.

### MRAP2 increases MC3R signaling

Previous studies have provided conflicting data regarding whether MRAP2 affects MC3R signaling (*2, 5, 18*). As our SiMPull data indicates that MRAP2 interacts with MC3R in a 1:1 stoichiometry, and there is no evidence that high concentrations of MRAP2 are required for its effects on MC3R, we performed our assays with equal concentrations of DNA. MC3R-induced increases in cAMP (assessed by Glosensor assays) were observed in cells expressing equal concentrations of MC3R and MRAP2 (Figure 3A-B). This effect was retained when transfecting as little as 25 ng of MC3R and MRAP2 (Figure S6) and therefore subsequent studies were performed using 25 ng of each plasmid to reduce overexpression artefacts. The endogenous antagonist AgRP was still able to inhibit MC3R activity in the presence of MRAP2 (Figure 3C). MRAP2 had no effect on MC3R cell surface expression when assessed using cell impermeable SNAP-647 labelling and fluorescence quantification or ELISA (Figure 3D-E).

**Figure 3.**
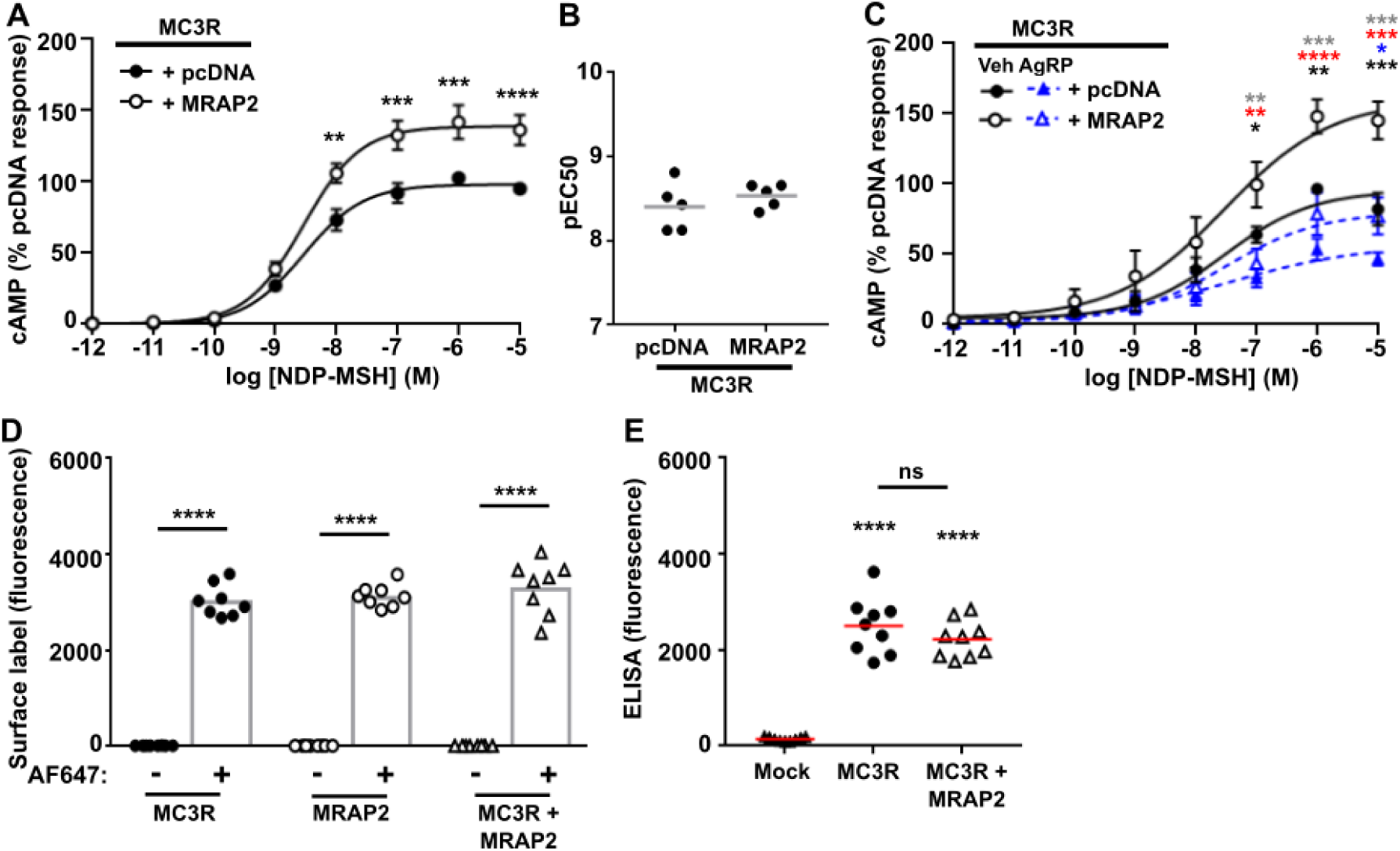
MRAP2 enhances MC3R signaling but has no effect on cell surface expression. (**A**) MC3R-induced cAMP responses measured by GloSensor in cells transfected with pcDNA or MRAP2. AUC was measured and responses expressed relative to the pcDNA maximal response. N=5. (**B**) pEC50 values from A. (**C**) Effect of the endogenous antagonist AgRP on MC3R-induced cAMP responses in cells transfected with pcDNA or MRAP2. N=6. Data shows mean±SEM in A and C and mean in B. Statistical analyses show pcDNA vs. MRAP2 with vehicle (black asterisks) or AgRP (blue), pcDNA vehicle vs. pcDNA AgRP (gray), MRAP2 vehicle vs. MRAP2 AgRP (red). (**D**) Surface labeling of cells transfected with SNAP-tagged MC3R, SNAP-MRAP2 or SNAP-MC3R with FLAG-CLIP-MRAP2 and labeled with SNAP-surface Alexa Fluor 647 (AF647). Fluorescence values were expressed relative to cells without the fluorescent label. There was no significant difference between MC3R or MRAP2 and combined MC3R+MRAP2. N=6. (**E**) Cell surface expression of MC3R assessed by ELISA in cells transfected with FLAG-CLIP-MC3R and SNAP-MRAP2 or pcDNA. Statistical analyses were performed by two-way ANOVA and Sidak’s for A and C and one-way ANOVA with Sidak’s for D-E. ****p<0.0001, ***p<0.001, **p<0.01, *p<0.05.

### Identification of residues required for MC3R and MRAP2 interactions

To understand how MRAP2 may interact with and facilitate MC3R signaling we used AlphaFold2 to predict structural homology models. We first predicted the structure of MC3R and MRAP2 in a 1:1 stoichiometry as the SiMPull data indicated this was the most common form of the heterodimer. Of the five predicted models, one had multiple side chain collisions that could not be reduced with model refinement, and large unstructured regions, and therefore was not further assessed (Figure S7). The other four models had a high confidence threshold, and predicted MRAP2 interacts with TM5-TM6 of MC3R, regions that are known to have an important role in receptor activation and G protein coupling to MC3R (*30*). The models predicted that MRAP2 may insert within the membrane in two orientations (i.e. an extracellular N-terminus in two models and intracellular in the other models), consistent with previous studies that indicated that MRAP2 may insert in this orientation (*27, 31*) (Figure S7). The model ranked with the highest confidence (Model 1) contained more structured regions than the other models, including a loop close to the ligand-binding pocket of MC3R and a helical structure in the juxtamembrane G protein-binding region (Figure 4A), similar to that observed in the MC2R-MRAP1 cryo-EM model (*32*).

**Figure 4.**
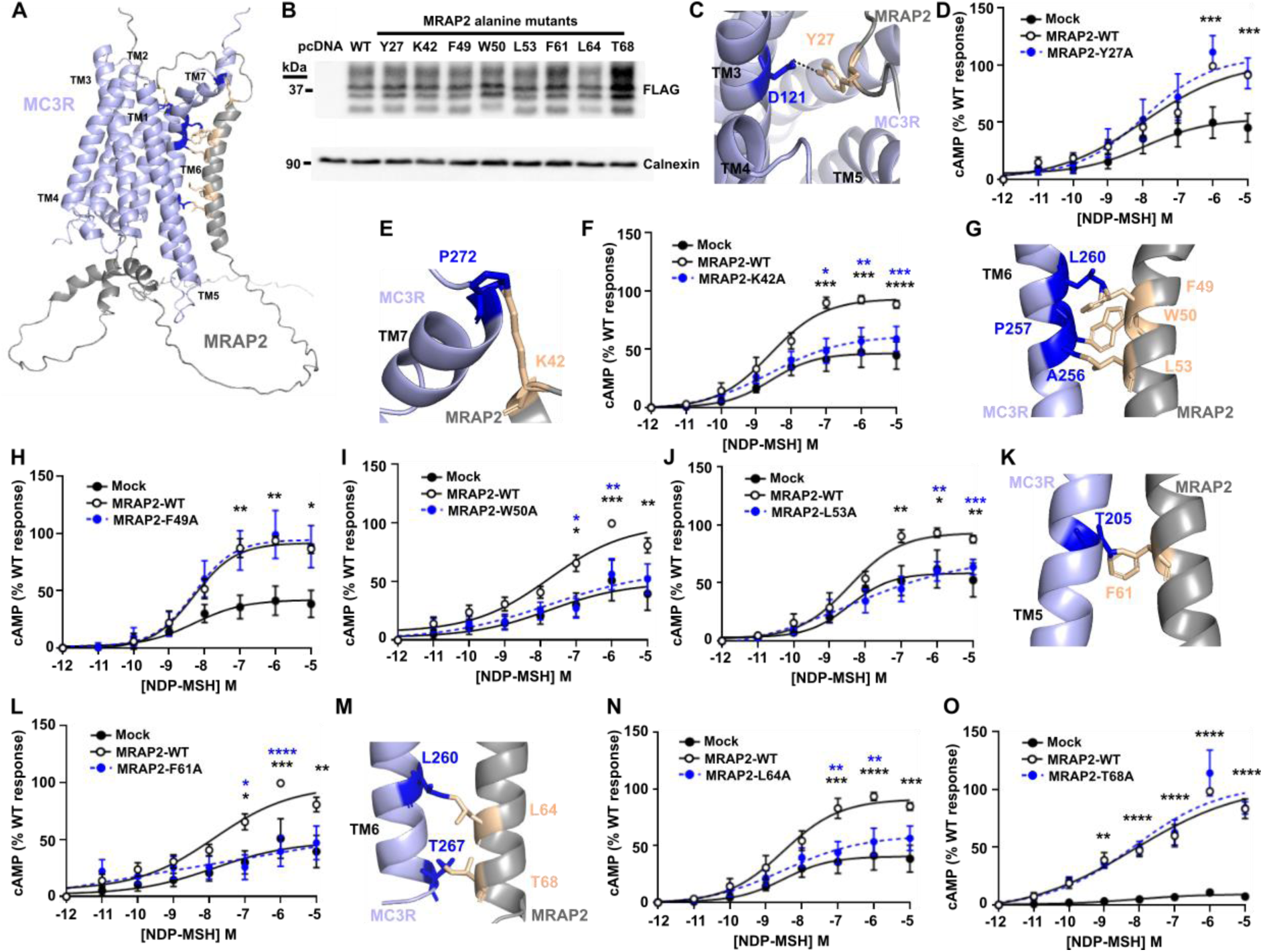
Prediction and assessment of MRAP2 residues that interact with MC3R. (**A**) Predicted structural model of MC3R and MRAP2 in a 1-to-1 configuration with residues tested in cAMP assays highlighted in orange. (**B**) Western blot showing expression of FLAG-MRAP2 alanine mutants. Calnexin was used as a housekeeper loading control. (**C**) Predicted contacts between MRAP2-Y27 and MC3R. The image shows the top of the structure with Tyr27 located close to the ligand-binding region of MC3R. (**D**) MC3R-induced cAMP responses for MRAP2-Y27A. N=6. (**E**) Predicted contacts between MRAP2-K42 and MC3R and (**F**) MC3R-induced cAMP responses for MRAP2-K42A. N=7. (**G**) Predicted contacts between MRAP2-F49, -W50, -L53 and MC3R. (**H-J**) MC3R-induced cAMP responses for MRAP2-F49 (N=7), -W50 (N=5), -L53 (N=6) and MC3R. (**K**) Predicted contacts between MRAP2-F61 and MC3R and (**L**) MC3R-induced cAMP responses for MRAP2-F61A. N=5. (**M**) Predicted contacts between MRAP2-L64 and -T68 and MC3R, and (**N-O**) MC3R-induced cAMP responses for MRAP2-L64A (N=7) and -T68A (N=4). Comparisons show MC3R variant and WT (blue) or MC3R variant and pcDNA (black). Statistical analyses were performed by two-way ANOVA with Sidak’s or Dunnett’s multiple-comparisons test. ****p<0.0001, ***p<0.001, **p<0.01, *p<0.05.

Models 1-4 were assessed to determine all possible contacts between MRAP2 and MC3R, which identified twenty-two possible interactions observed in at least one model (Table S3). We hypothesized that those residues identified in ≥3 models are more likely to be genuine contacts and therefore performed alanine mutagenesis of these residues in the FLAG-MRAP2 construct to determine whether they affect MC3R activity. We additionally assessed one residue (T68) located in the TM region close to these other residues that was predicted to form contacts in two models. Mutation of seven of these residues had no effect on the total protein and cell surface expression of MRAP2 or MC3R (Figure 4B, S8, Table S4). MRAP2-T68A significantly enhanced the total protein expression of MRAP2 (Figure 4B, Table S4) but did not affect the cell surface expression of either MRAP2 or MC3R (Figure S8). The Y27 residue is predicted to form contacts in all four models and lies in the ligand-binding region of MC3R in two models and the G protein docking region of two models (Figure 4C, S7). Mutation to alanine had no effect on MC3R-induced cAMP responses (Figure 4D). Seven residues in the TM region (K42, F49, W50, L53, F61, L64, T68) were predicted to form contacts with MC3R in multiple structural models (Table S3). Mutation of K42, W50, L53, F61 and L64 reduced MC3R-induced responses such that they were indistinguishable from MC3R responses in the absence of MRAP2. Alanine mutagenesis of the other residues had no effect on MC3R signaling (Figure 4E-O).

To investigate the MC3R-MRAP2 interaction in further detail we next mutated residues in MC3R that are predicted to interact with the five MRAP2 residues that impair MC3R-induced signaling (Table S3). Alanine mutagenesis was performed on three MC3R residues, Thr245 (TM6), Leu260 (TM6), Pro272 (TM7), and the effect on cAMP signaling first assessed in the absence of MRAP2. Mutagenesis of the three residues had no effect on MC3R cell surface expression (Figure 5A-B), and the Thr245Ala and Pro272Ala MC3R variants had no effect on agonist-induced responses in the absence of MRAP2. Leu260Ala reduced MC3R signaling and therefore this residue may have a role in MC3R activation that is distinct from MRAP2-induced effects (Figure 5C). Addition of MRAP2 did not further enhance MC3R-induced signaling by any residue above MRAP2-WT responses indicating that all three may contribute to MC3R-MRAP2 interactions (Figure 5D).

**Figure 5.**
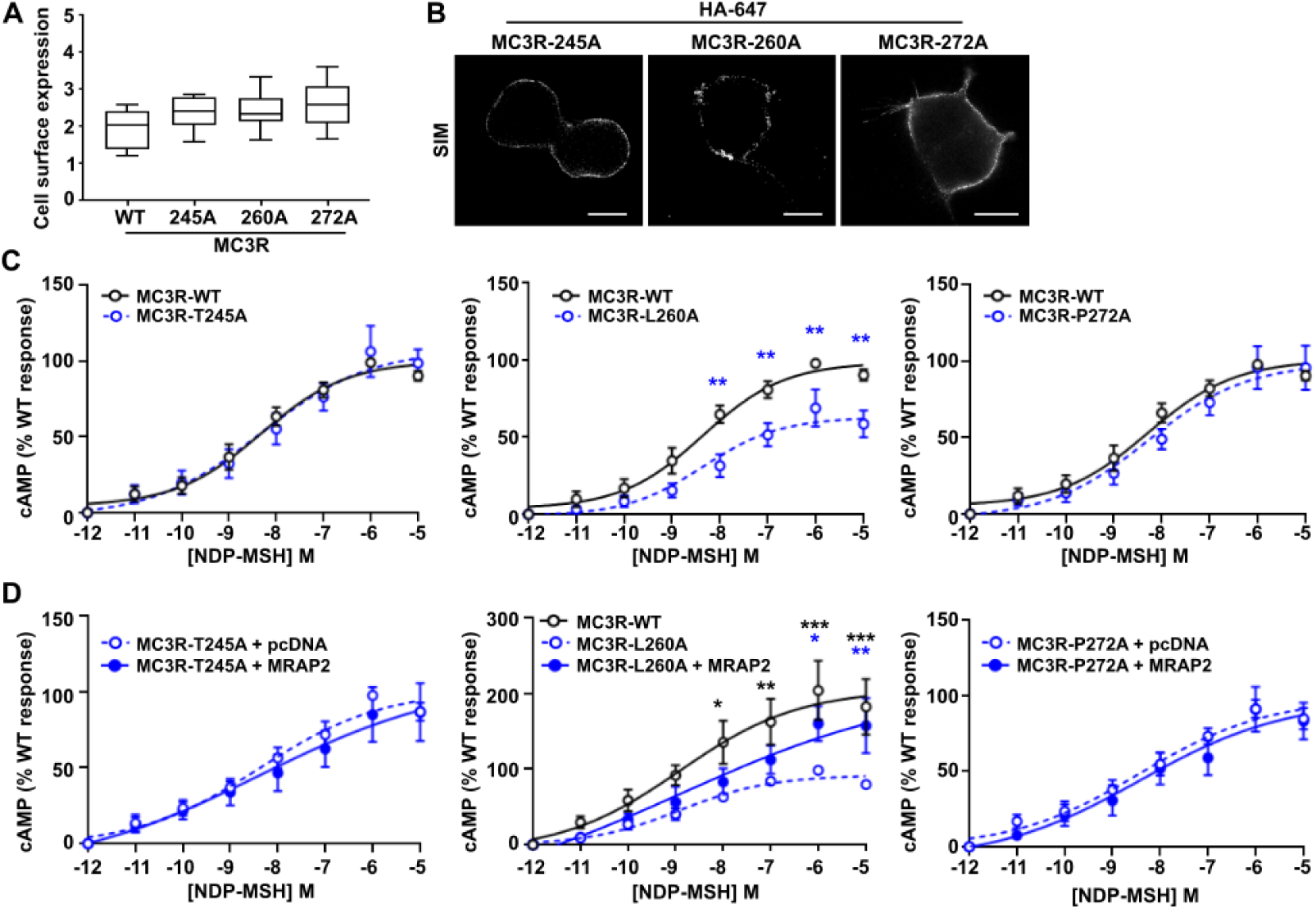
Assessment of MC3R residues that interact with MRAP2. (**A**) Fluorescent cell surface expression of MC3R alanine mutants compared to WT. N=6. (**B**) MC3R expression measured by SIM. Scale, 5 μm. (**C**) cAMP responses for MC3R-T245A, MC3R-T260A and MC3R-P272A compared to MC3R-WT. N=9 for all. (**D**) cAMP responses for MC3R-T245A, MC3R-T260A and MC3R-P272A with pcDNA or MRAP2. N=7 for all. Statistical analyses were performed by two-way ANOVA with Sidak’s or Dunnett’s multiple-comparisons test. Comparison between MC3R-alanine variant and WT (black) or MC3R-alanine variant with MRAP2 (blue). Statistical analyses were performed by two-way ANOVA with Sidak’s or Dunnett’s multiple-comparisons test. ***p<0.001, **p<0.01, *p<0.05.

As previous studies have suggested that dimeric MRAP2 interacts with MC3R, we also performed AlphaFold2 structural homology modelling with one MC3R and two MRAP2 residues. These models did not predict MC3R interactions with dimeric MRAP2, and instead predicted that the two MRAP2 residues may interact in two distinct sites on MC3R (Figure S7). As this correlated with the SiMPull data that indicated binding of monomeric MRAP2 is preferential, we did not investigate these models in further detail.

### MRAP2 increases MC3R internalization

Previous studies have shown that MRAP2 enhances GPCR signaling by impairing β-arrestin recruitment and consequently reducing receptor internalization (*20, 22, 23*). To determine whether MRAP2 uses a similar mechanism to enhance MC3R signaling, bystander BRET assays were performed measuring proximity between Nluc-tagged β-arrestin-2 and Venus-tagged Kras, a marker of the plasma membrane, in the presence of MC3R. BRET was enhanced in a concentration-dependent manner in all cells although responses were significantly reduced in MRAP2 transfected cells when compared to control cells (Figure 6A). This suggests that MRAP2 impairs MC3R-mediated β-arrestin-2 recruitment and to further investigate this we assessed β-arrestin-2-YFP expression in MC3R expressing cells by SIM imaging. Under basal conditions β-arrestin-2 is distributed across the cytoplasm in cells expressing MRAP2 or pcDNA control (Figure 6B). In the presence of agonist, β-arrestin-2 is recruited to the plasma membrane rapidly in control cells. In cells expressing MRAP2, β-arrestin-2-YFP forms punctate structures, but plasma membrane recruitment is only apparent following 20 minutes exposure to agonist (Figure 6B).

**Figure 6.**
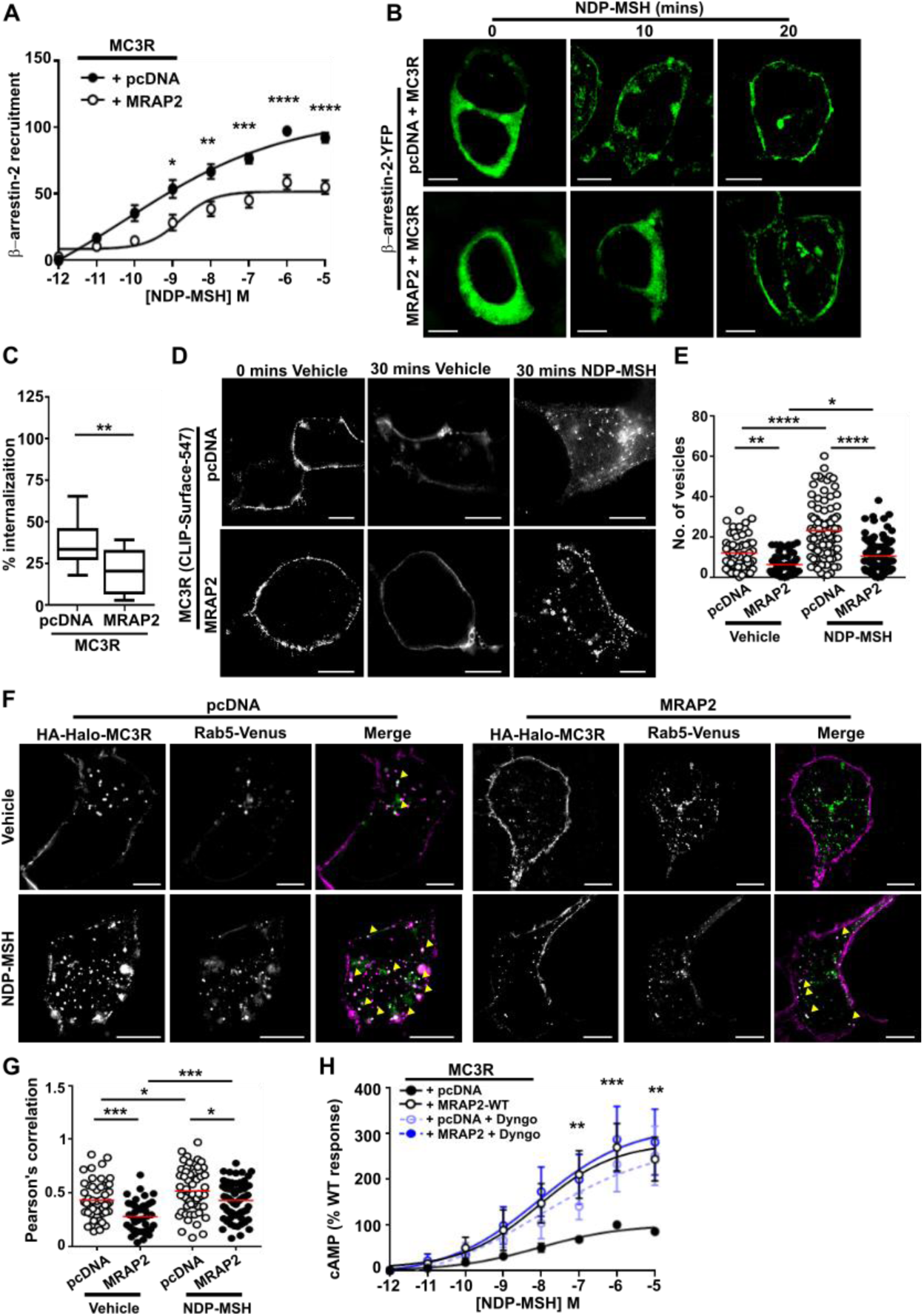
MRAP2 reduces β-arrestin recruitment and impairs receptor internalization. MC3R-induced membrane recruitment of β-arrestin-2 with pcDNA and MRAP2 measured by (**A**) BRET (N=8) and (**B**) SIM. Scale, 5 μm. (**C**) Percentage internalization of SNAP-MC3R following exposure to NDP-MSH for 30 minutes in cells transfected with pcDNA or MRAP2. N=12. (**D**) Agonist-induced internalization assessed by SIM imaging of BC-DY547-labeled MC3R in the presence of pcDNA or MRAP2. Scale, 5 μm. (**E**) Quantification of the number of internalized vesicles in cells exposed to vehicle or NDP-MSH for 30 minutes. N=56-57 cells (vehicle) and N=91-93 cells (agonist) from seven independent transfections for each group. (**F**) SIM imaging of MC3R and Rab5 in the presence of pcDNA or MRAP2. N=41-60 cells from five independent transfections for each group. Scale, 5 μm. (**G**) Correlation between MC3R and Rab5 in SIM images assessed by Pearson’s coefficient. (**H**) MC3R-induced cAMP responses in the presence of pcDNA or MRAP2 ± Dyngo (N=6) Statistical analyses were performed by two-way ANOVA with Sidak’s or Dunnett’s multiple-comparisons test in A, H, one-way ANOVA with Sidak’s test in E and G, and unpaired t-test in C. ****p<0.0001, ***p<0.001, **p<0.01, *p<0.05.

As MRAP2 impaired MC3R-induced β-arrestin-2 recruitment to the plasma membrane it was hypothesized that MRAP2 would reduce receptor internalization. To assess this, cells were transfected with CLIP-MC3R, labeled with cell impermeable BC-DY547, then the amount of surface labelled receptor quantified following exposure to vehicle or agonist for 30 minutes in 96-well plates. Surface labeling of MC3R was reduced following exposure to ligand, consistent with agonist-induced internalization of the receptor. When the percentage difference was quantified, there was a significantly greater internalization in control cells than MRAP2 expressing cells (Figure 6C). To assess internalization in more detail cells were transfected with HA-HALO-MC3R and incubated with an HA antibody and either vehicle or NDP-MSH for 30 minutes, then imaged by SIM. Endocytosis of fluorescently-labeled MC3R was apparent in control and MRAP2 expressing cells exposed to ligand, although there appeared to be more internalized receptor in cells transfected with MC3R without MRAP2 (Figure 6D). When the number of vesicles was quantified, MRAP2 expressing cells had fewer vesicles in both vehicle and NDP-MSH treated cells when compared to cells expressing empty vector, indicating that both constitutive and agonist-driven MC3R internalization is reduced by MRAP2 (Figure 6E). Consistent with reduced internalization, there was significantly less colocalization between the early endosome marker Rab5 and MC3R in MRAP2 expressing cells when assessed by SIM (Figure 6F-G) in both vehicle and agonist exposed cells. Thus, MRAP2 impairs both constitutive and agonist-driven MC3R internalization.

These studies suggest that MRAP2 may enhance MC3R signaling by retaining the receptor at the cell surface due to reduced internalization. To assess whether blocking MC3R internalization results in an increase in receptor signaling, cells were pre-treated with Dyngo-4a, which impairs clathrin-mediated endocytosis (Figure S9), then signaling assessed by cAMP Glosensor assays. Impairment of internalization enhanced MC3R-induced cAMP responses in cells expressing pcDNA, such that these were no longer different to MRAP2 responses (Figure 6H). Pre-treatment of MRAP2 expressing cells with Dyngo-4a had no effect on MRAP2 responses. This suggests impaired receptor internalization is one mechanism by which MRAP2 enhances GPCR signaling.

### Obesity-associated variants in MRAP2 impair MC3R function

Previous studies have identified associations between MRAP2 genetic variants and obesity, hypertension and diabetes (*5, 6*). It is possible that these variants may affect the function of other GPCRs that MRAP2 associates with, and we therefore examined MC3R function in HEK293 cells expressing twelve different MRAP2 human variants. The twelve MRAP2 variants were selected based on their predicted location (Figure 7A) either in the N-terminal ligand-binding region (G31V, P32L), transmembrane domain (F62C), C-terminal unstructured region (N88Y, V91A) or C-terminal helical structure within the G protein binding region (R113G, S114A, L115V, N121S, R125C, H133Y, T193A). The variants had no significant effect on MC3R expression at the plasma membrane (Figure S10). The G31V and P32L variants were not predicted to affect interactions with MC3R in the AlphaFold2 models (Table S5) and had no effect on MC3R-mediated cAMP responses (Figure 7B). MRAP2-F62 forms backbone interactions with other residues within the MRAP2 transmembrane helix (Table S5). The variant MRAP2-F62C significantly impaired MC3R-mediated cAMP responses such that they are not significantly different to cells expressing pcDNA. MRAP2-N88Y also significantly reduced MC3R-mediated cAMP responses (Figure 7C). The R113G and S114A variants were predicted to lose contacts with adjacent MC3R and MRAP2 residues, respectively (Figure 7D-E, Table S5), and significantly impaired MC3R-induced cAMP signaling, as did the neighbouring L115V variant (Figure 7F-G). Similarly, three other variants within the MRAP2 C-terminus (N121S, R125C, T193A) also significantly reduced MC3R activity (Figure 7H-I).

**Figure 7.**
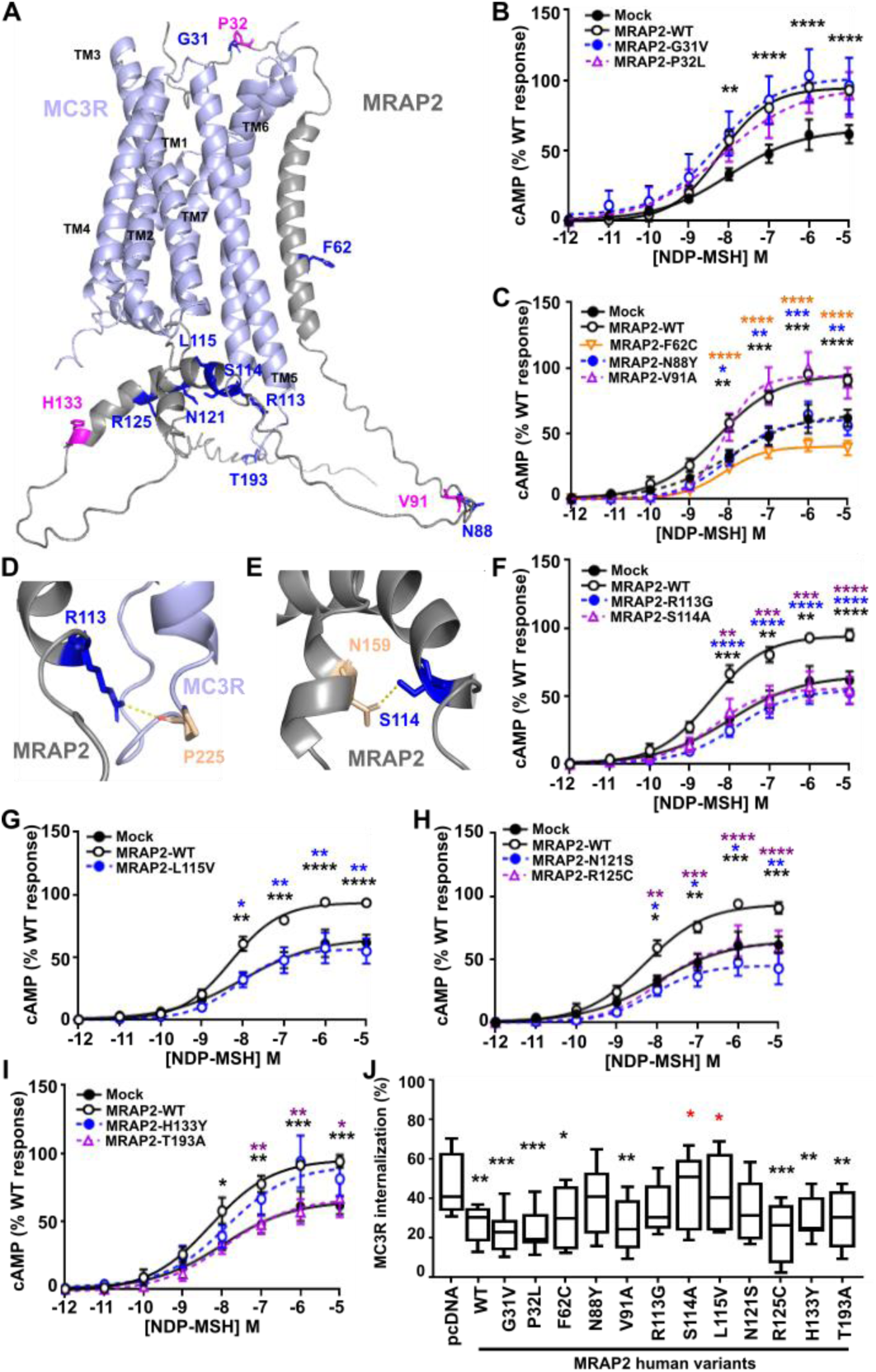
Effect of human MRAP2 variants on MC3R signaling and internalization. (**A**) Predicted structural model of MC3R and MRAP2 in a 1-to-1 configuration with residues mutated in overweight or obese individuals. Residues highlighted in pink have been identified only in normal weight individuals. (**B**) MC3R-induced cAMP responses for MRAP2-G31V (N=7) and -P32L (N=6). (**C**) MC3R-induced cAMP responses for F62C (N=7), N88Y (N=7) and V91A (N=5). (**D-E**) Predicted contacts between MRAP2-R113, -S114 and MC3R. (**F-I**) MC3R-induced cAMP responses for (F) MRAP2-R113G, -S114A, (G) L115V, (H) N121S, R125C, (I) H133Y, T193A. N=7 for F-H, N=6 for I. (**J**) MC3R-induced internalization in cells expressing pcDNA, MRAP2 wild-type or the twelve MRAP2 variants. Statistical analyses were performed by two-way ANOVA with Sidak’s or Dunnett’s multiple-comparisons test. Asterisks compare MRAP2 wild-type to: pcDNA in black, variants according to their labeling in blue, orange or purple in A-I. In panel J asterisks compare variants to pcDNA in black and to wild-type in red. ****p<0.0001. ***p<0.001, **p<0.01, *p<0.05.

The effect of the variants on MC3R internalization was examined by labeling cells with SNAP-surface-647 following exposure to vehicle or agonist for 30 minutes and quantifying the surface labeling. Fluorescence was reduced in all cells, consistent with agonist-induced internalization. Most variants had a similar effect on MC3R internalization as the wild-type MRAP2 protein (i.e. significantly decreased internalization compared to pcDNA) (Figure 7J). Two MRAP2 variants, S114A and L115V, had internalization levels that were significantly different to MRAP2 wild-type and instead had similar internalization to that observed in cells transfected with pcDNA, indicating that these variants impair the effects of MRAP2 on internalization (Figure 7J). Three variants, N88S, R113G and N121S, had an intermediate profile which was not significantly different to pcDNA or MRAP2 wild-type, indicating that these may partially impair MRAP2’s effect on internalization.

## Discussion

Our studies have shown that MRAP2 interacts with MC3R to enhance signaling and expands the repertoire of receptors that have been robustly demonstrated to interact with MRAP2. While MRAP2 has been described to interact with >40 GPCRs, signaling data has not been provided for most receptors, and often the data is not replicable between different groups (*2, 3, 5, 13*). In contrast, our data provides multiple lines of evidence demonstrating that MRAP2 facilitates MC3R signaling. Firstly, we have shown that MC3R and MRAP2 are co-expressed in human neurons that regulate energy homeostasis and food intake, and that the proportion of neurons that co-express MC3R and MRAP2 is similar to MC4R and MRAP2 co-expression, which are widely accepted to interact (Figure 1, S1, Table S1). Secondly, the two proteins interact at the single-molecule level, and MRAP2 enhances signaling at low levels of expression (Figure 2-3). Thirdly, disruption of putative interacting residues impairs MRAP2-mediated signaling (Figure 4-5). Fourthly, MRAP2 uses similar mechanisms to those described for other receptors to impair β-arrestin recruitment (Figure 6). Finally, human variants in the MRAP2 transmembrane domain and C-terminus implicated in receptor interactions impair signaling and affect internalization (Figure 7). Therefore, we are confident that MC3R and MRAP2 form heterodimers that contribute to MC3R function.

Several groups have shown that MRAP2 may form dimers at the cell surface (*2, 18, 28*), and this has led to the assumption that MRAP2 exists in a dimeric form at the membrane which is necessary for its function (*7, 33*). However, studies of MRAP2 dimerization examined the protein in isolation and therefore the effect of co-transfected GPCRs and heterodimer stoichiometry on MRAP2 were not established. Our SiMPull experiments (Figure 2) show that MRAP2 can form dimers at the cell surface, consistent with previous studies, although, monomeric MRAP2 is more prevalent. Moreover, when co-expressed with receptor, most complexes comprise one MC3R molecule interacting with monomeric MRAP2. Consistent with this stoichiometry, our structural homology models similarly predicted binding by monomeric MRAP2, and published structures of MC2R and MRAP1 also have a 1:1 stoichiometry (*32*). It is possible that in an environment in which GPCR expression is low, MRAP2 may form dimers at the membrane, but when co-expressed with GPCRs it favours heterodimerization with the receptor in a monomeric form. However, our SiMPull analyses demonstrated a sizable proportion of MRAP2 is in a monomeric form at cell surfaces when the cells were transfected only with HALO-MRAP2, and therefore it is likely that there are both monomers and dimers at the cell surface, although our choice of detergent could have impacted these quantities. Our cell surface labeling strategy with membrane impermeable dyes would also not be able to detect MRAP2 inserted in a C-terminal out orientation, and therefore we cannot discount that these dimers may also form. Examination of additional complexes will be required to determine whether this 1:1 stoichiometry is important for other MRAP2-GPCR interactions. As our studies, and those of a recent preprint (*22*), have shown that overexpression of MRAP2 is unnecessary, future studies should also assess the 1-to-1 stoichiometry.

We identified five residues in MRAP2 that may contribute to receptor interactions and/or facilitate signalling. These residues are all located in the transmembrane helix, a region that has previously been shown to be important for potentiation of GHSR signalling (*20*) by MRAP2. The transmembrane region is also important for MRAP1 interactions with MC2R (*34*), and it is likely that there is a shared mechanism by which these accessory proteins facilitate GPCR signaling. Cryo-EM structures of MC2R with MRAP1 demonstrate that the accessory protein interacts with TM5 and TM6 of the receptor (*32*). Our homology models of MC3R and MRAP2 predict interactions with TM5 or TM6 of the receptor, and alanine mutagenesis of three residues within TM6 impaired MRAP2’s ability to facilitate MC3R signalling. MC3R conforms to common class A G protein coupling mechanisms whereby outward movement of TM6 allows formation of a large cytoplasmic cavity between TM5-TM7 that can accommodate G protein binding (*30*). Similar activation mechanisms have been described for MC2R (*32*) and MC4R (*35*), and we propose that MRAP2-TMD interactions with TM5-TM6 of GPCRs, allows the receptor to adopt a structural conformation that is more readily activated and/or allows G proteins to couple more efficiently. Such facilitation of a ‘partially preactivated state’ that can be more readily activated has been described for the RAMP2 accessory protein’s ability to potentiate signalling by the parathyroid hormone type-1 receptor (PTH1R) (*36*). Interactions between TM4 and TM5 of PTH1R and the RAMP2 transmembrane domain are important for establishing this preactivated state.

Consistent with previous studies of other GPCRs (*20, 22, 23*) we also found that MRAP2 impairs β-arrestin recruitment. Although the mechanism by which MRAP2 impairs β-arrestin binding is unknown, the AlphaFold2 models suggest that the intracellular MRAP2 α-helix may sterically block the β-arrestin binding site which is likely to involve the intracellular ends of TM5 and TM6 (*37*). However, such a mechanism would also be expected to impair G protein coupling suggesting that the cytoplasmic α-helix may undergo conformational changes following receptor activation. Further studies of the structure of the MRAP2 cytoplasmic region could provide insights into these mechanisms. Reduced β-arrestin recruitment and the consequent impairment in receptor internalization could explain some of the effects of MRAP2 on GPCR activation. Consistent with this, blocking MC3R endocytosis using Dyngo-4a enhanced receptor signaling and previous studies have shown that AgRP inhibits MC3R activity, at least in part, by enhancing recruitment of β-arrestin and promoting receptor endocytosis (*38*). However, while we showed multiple MRAP2 variants impaired MC3R cAMP signaling, not all affected receptor trafficking, and it is possible that other mechanisms exist that allow MRAP2 to promote receptor signaling.

Previous studies of MRAP2 in human populations identified >25 variants associated with obesity (*5–7, 14, 39*). One group also reported hyperglycaemia and hypertension occurred more commonly in individuals with MRAP2 mutations than in those with MC4R and suggested that MRAP2 variants may affect signaling by other GPCRs (*6*). This was based on the finding that the variants did not all impair MC4R function, and the phenotype was dissimilar to that in individuals with MC4R mutations. Our studies show that eight MRAP2 variants impair MC3R-mediated cAMP activity, while three variants (P32L, V91A, H133Y) found exclusively in individuals with normal weight (*6*) had no effect on signaling or trafficking. Whether inhibition of MC3R contributes to any of the clinical findings identified in individuals with MRAP2 variants is unknown. Mice with deletion of *Mc3r* have a high ratio of fat-to-lean mass but are not markedly obese unless fed a high fat diet, and heterozygous mice have normal weight (*40–42*). However, mice depleted of both *Mc3r* and *Mc4r* are significantly heavier than *Mc4r^-/-^* mice (*40*), suggesting MC3R can contribute to weight gain. In contrast, depletion of *Mc3r* from AgRP neurons causes an anorexia and starvation phenotype, consistent with its known orexigenic role in these neurons (*43*). In humans, rare inactivating MC3R variants have been associated with obesity, but these findings are inconsistent (*44*). Recently, several functionally inactivating MC3R heterozygous mutations have been linked to childhood growth and timing of puberty with normal weight (*45*). One homozygous individual had also been overweight/obese since childhood and had type-2 diabetes and hypertension (*45*). Therefore, further studies of individuals with MRAP2 or MC3R variants are required to better understand how inactivating variants contribute to disease.

We also showed that MRAP2 variants can affect pathways other than the cAMP pathway. Five variants located in the intracellular domain that impaired cAMP signaling also reduced internalization. Although we know little about the MRAP2 C terminus, structural homology models suggest part of this region may form an α-helical structure that lies within the MC3R cleft in which G proteins and β-arrestin bind. Several of the variants (R113G, S114A, L115V, N121S) that affect both signaling and trafficking are present in this structure and we hypothesise that these variants disrupt the ability of the MC3R to engage with G proteins, resulting in impaired signaling. It will be important to investigate whether MRAP2 variants affect multiple aspects of GPCR signaling, as studies of MC4R have demonstrated inactivating mutations that contribute to obesity may not affect canonical signaling, but can affect internalization, homodimerization or other G protein pathways (*46*). Moreover, MRAP2 variants should be tested to determine whether they affect signaling by other GPCRs.

In summary, we have shown that MRAP2 directly binds to MC3R to enhance Gs-mediated signaling and impair β-arrestin recruitment. Our mutagenesis studies and examination of human genetic variants demonstrated that the MRAP2 transmembrane domain and a putative C-terminal helix play an important role in facilitating MRAP2-mediated enhancement of MC3R activity and may have applicability to other GPCRs. Novel therapies that disrupt or enhance these sites could have important implications for treating disorders of food intake including obesity and anorexia.

## Materials & Methods

### Plasmid constructs and compounds

A full list of plasmids with their source can be found in Table S2. For single molecule pull-down experiments, constructs were generated with an N-terminal signal peptide from rat mGluR2 (*25*), followed by affinity tags (HA or FLAG), self-labeling protein tags capable of conjugation to organic dyes (SNAP, CLIP, or Halo), and human MC3R and MRAP2. Cloning into the pRK5 vector was performed using reagents from Promega and oligonucleotides from Sigma to generate the following plasmids: ss-HA-Halo-MC3R, ss-HA-SNAP-MC3R, ss-FLAG-CLIP-MC3R, ss-HA-Halo-MRAP2, ss-HA-SNAP-MRAP2, ss-FLAG-CLIP-MRAP2, ss-HA-HALO-MC4R, ss-FLAG-CLIP-SSTR3. The MRAP2 variants were introduced into a MRAP2-3xFLAG plasmid by site-directed mutagenesis using the Quikchange Lightning Kit (Agilent Technologies) and oligonucleotides from Sigma. All plasmids were sequenced verified by Source Bioscience. NDP-MSH (Cambridge Bioscience) was used at a concentration of 10 µM, unless otherwise stated, Dyngo-4a (Abcam) was used at a concentration of 30 µM with cells pre-incubated for 30 minutes prior to experiments, AgRP (Bio-Techne) was used at 0.1 µM.

### Cell culture and transfection

Adherent HEK293 cells were purchased from Agilent Technologies and were maintained in DMEM-Glutamax media (Merck) with 10% calf serum (Merck) at 37°C, 5% CO_2_. Cells were routinely screened to ensure they were mycoplasma-free using the TransDetect Luciferase Mycoplasma Detection kit (Generon). Expression constructs were transiently transfected into cells using Lipofectamine 2000 (LifeTechnologies), following manufacturer’s instructions.

### Transcript expression analysis

To assess the extent of co-expression of *MRAP2* with *MC3R* and *MC4R* separately we utilised HYPOMAP: a spatio-cellular atlas of the human hypothalamus (*24*). Log-normalised gene expression for MRAP2 and MC3R was visualized in the spatial transcriptomics dataset. To highlight co-expression, spots which expressed both *MC3R* and *MRAP2* transcripts were highlighted. Using the single nucleus RNA-sequencing dataset, we calculated the percentage of neurons which expressed MRAP2 across the whole hypothalamus dataset, as well as the percentage of *MC3R*-positive neurons that co-expressed *MRAP2*, and the percentage of *MC4R*-positive neurons that co-expressed *MRAP2*. Co-expression was also measured on a cluster-by-cluster basis, at the highest resolution of clustering. To highlight co-expression in the snRNAseq dataset, cells which expressed MRAP2 and MC3R transcripts, or MRAP2 and MC4R transcripts were highlighted in the UMAP plots. Analysis and plots were performed using R and ggplot2.

### NanoBiT assays

NanoBiT assays were performed using methods adapted from previous studies (*47*). MRAP2 and MC3R were cloned into the LgBiT-C and SmBiT-C plasmids (purchased from Promega). HEK293 cells were seeded at 10,000 cells/well in 96-well plates and transfected the same day with 100ng (or as specified in the relevant figure legend) LgBiT and SmBiT plasmids. Following 48-hours, media was changed to FluoroBrite DMEM phenol red-free media (Gibco) with 10% calf serum (FluoroBrite complete media) with 40 μL Nano-Glo substrate (Promega) and luminescence baseline signals read on a Glomax (Promega) plate reader at 37 °C. Data was normalized to luminescence values in the negative control (MC3R-SmC and LgC-Empty).

### Single molecule pull-down (SiMPull)

Cells were seeded in 12-well plates and transfected with 300 ng of Halo-tagged plasmids and 600ng of CLIP-tagged plasmids. After 24 hours, cells were washed with extracellular solution (comprising 135 mM NaCl (Sigma), 5.4 mM KCl (Sigma), 10 mM HEPES (Gibco), 2 mM CaCl_2_ (VWR Chemicals); 1 mM MgCl_2_ (Sigma), pH 7.4), then labelled with 2 μM of cell-membrane impermeable dyes (CLIP-surface 547 (BC-DY547, NEB) for FLAG-CLIP tagged plasmids, or CA-sulfo646 for HA-Halo tagged plasmids) in extracellular solution for 45 min at 37 °C. Cells were washed with extracellular solution, harvested in 1x Ca^2+^- and Mg^2+^-free PBS, then cell pellets lysed in buffer (Tris pH8, NaCl, EDTA (all from Sigma)) containing 0.5% Lauryl Maltose Neopentyl Glycol/ 0.05% Cholesteryl Hemisuccinate (LMNG-CHS) (Anatrace) and protease inhibitor (Roche). Microflow chambers were prepared by passivating a glass coverslip and quartz slide with mPEG-SVA and biotinylated PEG (MW = 5000, 50:1 molar ratio, Laysan Bio), as previously described (*25, 48*). Prior to each experiment a chamber was incubated with 0.2 mg/ml NeutrAvidin (Fisher Scientific UK) for 2 min, washed in T50 buffer (50 mM NaCl, 10 mM Tris), then incubated with 10 nM biotinylated anti-HA antibody (ab26228, Abcam, RRID:AB_449023) in T50 buffer (50 mM NaCl, 10 mM Tris) for 30 minutes. Fresh cell lysates were mixed with dilution buffer (1:10 lysis working solution with extracellular solution) and added to the flow chamber until a suitable single molecule spot density was obtained. Chambers were washed with dilution buffer to remove unbound receptor, then single molecule movies obtained as previously described (*25*) using a 100x objective (NA 1.49) on an inverted microscope (Olympus IX83) with total internal reflection (TIR) mode at 20 Hz with 50 ms exposure time with two sCMOS camera (Hamamatsu ORCA-Flash4v3.0). Samples were excited with 561 nm and 640 nm lasers to excite BC-DY547 and CA-Sulfo-646, respectively. Single molecule movies were recorded sequentially at 640 nm, then 561 nm until most molecules were bleached in the field. Images were analyzed using a custom-built LabVIEW program (*49*). Each movie was concatenated using MatLab (R2022a), then loaded on LabVIEW to visualize each channel for co-localized molecules. The fluorescence trace of each molecule was inspected manually and bleaching steps aligned. Data were plotted using GraphPad Prism.

### Three-dimensional modeling of MRAP2 and MC3R

Modeling of MC3R and MRAP2 was performed by AlphaFold2 using the ColabFold v1.5.2-patch in Google Co-laboratory (*50*) and visualized using Pymol. FASTA sequences were obtained from NCBI. Five models were predicted and ranked based on predicted local distance difference test (pLDDT).

### Assessment of cell surface expression and internalization

For assessment of MC3R surface expression, cells were transfected with 100ng HA-HALO-MC3R and 100 ng pcDNA or FLAG-MRAP2 (wild-type of mutant) and cells fixed 48-hours later in 4% PFA (Fisher Scientific UK) in PBS, then labelled with 1:1000 anti-HA mouse monoclonal antibody (BioLegend Cat#901514, RRID:AB_2565336) followed by Alexa Fluor 647 donkey anti-mouse secondary antibody (abcam Cat# ab181292, RRID:AB_3351687). Cells were washed, then fluorescence read on a Glomax plate reader. Data was normalized to that observed in cells transfected with pcDNA, set as 1 and not shown on the graph.

For assessment of MC3R internalization in the presence of MRAP2 mutants, HEK293 cells were seeded at 10,000 cells/well in 96-well plates and transfected the same day with 100 ng HA-SNAP-MC3R and 100 ng pcDNA or MRAP2 (WT or variants). Forty-eight hours later, cells were exposed to 10 μM NDP-MSH or vehicle for 30 minutes, then labelled with SNAP-surface-647.

### Western blot analysis

For MRAP2 expression studies, either 3xFLAG-MRAP2-WT or 3xFLAG-MRAP2-mutants were transfected at 1 µg per well in a 6-well plate. Cells were lysed 48-hours later in NP40 buffer and western blot analysis performed as described (*51*). Blots were blocked in 5% marvel/TBS-T, then probed with anti-FLAG (M2 antibody, Sigma) and anti-calnexin (Millipore, Cat# AB2301, RRID:AB_10948000) antibodies. Blots were visualized using the Immuno-Star WesternC kit (BioRad) on a BioRad Chemidoc XRS+ system. Densitometry was performed using ImageJ (NIH), and protein quantities normalized to calnexin.

### Bioluminescence resonance energy transfer (BRET)

NanoBRET assays were performed using methods adapted from previous studies (*52*). HEK293 cells were seeded at 10,000 cells/well in 96-well plates and transfected the same day with 50 ng Nluc-Arr2, 500ng Venus-Kras, 100ng HA-Halo-MC3R and 100 ng pcDNA or FLAG-MRAP2. Forty-eight hours later, media was removed and replaced with Fluorobrite complete medium. Nano-Glo reagent was then added at a 1:100 dilution and BRET measurements recorded using a Promega GloMax microplate reader at donor wavelength 475-30 and acceptor wavelength 535-30 at 37 °C. The BRET ratio (acceptor/donor) was calculated for each time point. Four baseline recordings were made, then agonist added at 8 minutes and recordings made for a further ∼40 minutes. The average baseline value recorded prior to agonist stimulation was subtracted from the experimental BRET signal. All responses were then normalized to that treated with vehicle to obtain the normalized BRET ratio. AUC was calculated in GraphPad Prism and these values used to plot concentration-response curves with a 4-parameter sigmoidal fit.

### Glosensor cAMP assays

HEK293 cells were plated in 6-well plates and transfected with 200 ng pGloSensor-20F plasmid, and equal amounts of MC3R and MRAP2 (25-500 ng for transfection tests, and 25 ng for all other studies). Forty-eight hours later, cells were seeded in 96-well plates in FluoroBrite complete medium. Cells were incubated for at least 4 hours, then media changed to 100 µL of equilibration media consisting of Ca^2+^- and Mg^2+^-free HBSS containing 2% (v/v) dilution of the GloSensor cAMP Reagent stock solution (Promega). Cells were incubated for 2 h at 37°C. Basal luminescence was read on a Glomax plate reader for 8 min, then agonist added and plates read for a further 30 minutes. For FLAG-CLIP-SSTR3 studies, cells were preincubated with 10 μM forskolin for 5 minutes to elevate cAMP levels, then assays performed as described for MC3R with somatostatin-14 (Sigma) added as the agonist. Data was plotted in GraphPad Prism, area under the curve calculated and these values used to plot concentration-response curves with a 4-parameter sigmoidal fit.

### Structured illuminated microscopy (SIM)

Cells were plated on 24 mm coverslips (VWR) and transfected with 500ng of each plasmid 36-hours prior to experiments. For studies of cell surface expression, cells were fixed with 4% PFA in PBS and exposed to 1:1000 anti-HA mouse monoclonal antibody (BioLegend Cat#901514, RRID:AB_2565336) or 1:1000 anti-FLAG mouse monoclonal antibody (M2, Sigma-Aldrich), followed by Alexa Fluor 647 goat anti-mouse (Cell Signaling Technology Cat# 4410, RRID:AB_1904023). For MC3R and MRAP2 colocalization studies, the anti-HA rabbit primary antibody (ab26228, Abcam) was used with the anti-FLAG antibody, followed by Alexa Fluor 647 goat anti-mouse and Alexa Fluor 488 goat anti-rabbit (Cell Signaling Technology Cat# 4412, RRID:AB_1904025). For studies with Rab5-Venus, cells were exposed to 1:1000 anti-HA mouse monoclonal antibody (BioLegend Cat#901514, RRID:AB_2565336) with either vehicle or 10 μM NDP-MSH for 30 minutes. Cells were fixed, permeabilized and exposed to the Alexa Fluor 647 secondary antibody. Samples were imaged on a Nikon N-SIM system (Ti-2 stand, Cairn TwinCam with 2 × Hamamatsu Flash 4 sCMOS cameras, Nikon laser bed 488 and 647 nm excitation lasers, Nikon 100 × 1.49 NA TIRF Apo oil objective). SIM data was reconstructed using NIS-Elements (v. 5.21.03) slice reconstruction. Colocalization and Pearson’s correlation coefficient was measured using the ImageJ plugin JACoP.

### Statistical analysis

Statistical tests used for each experiment are indicated in the legends of each figure and the number of experimental replicates denoted by N. Data was plotted and statistical analyses performed in Graphpad Prism 7. Normality tests (Shapiro-Wilk or D’Agostino-Pearson) were performed on all datasets to determine whether parametric or non-parametric statistical tests were appropriate. A p value of <0.05 was considered statistically significant.

## Author contributions

Conceptualization: CMG

Methodology: GSHY, JLev, CMG

Investigation: AJ, RAW, Jlee, GD, JAT, CMG

Materials: JB, GKCD, GY, Jlev

Writing – original draft: CMG

Writing – review and editing: All authors

## Data and materials availability

All data needed to evaluate the conclusions in the paper are present in the paper and/or the Supplementary Materials. Plasmid constructs developed for this manuscript (see Table S2) will be made available upon request. Plasmid constructs obtained from other researchers are detailed in Table S2 and may be subject to Material Transfer Agreements. Please contact the corresponding author of this manuscript, or the named source for details.

