## Supplementary Figures for "The MRAP2 accessory protein directly interacts with melanocortin-3 receptor to enhance signaling"

**Supplementary Appendix**

**Figure S1 Expression of MRAP2 and MC3R in the human hypothalamus measured by spatial transcriptomics**

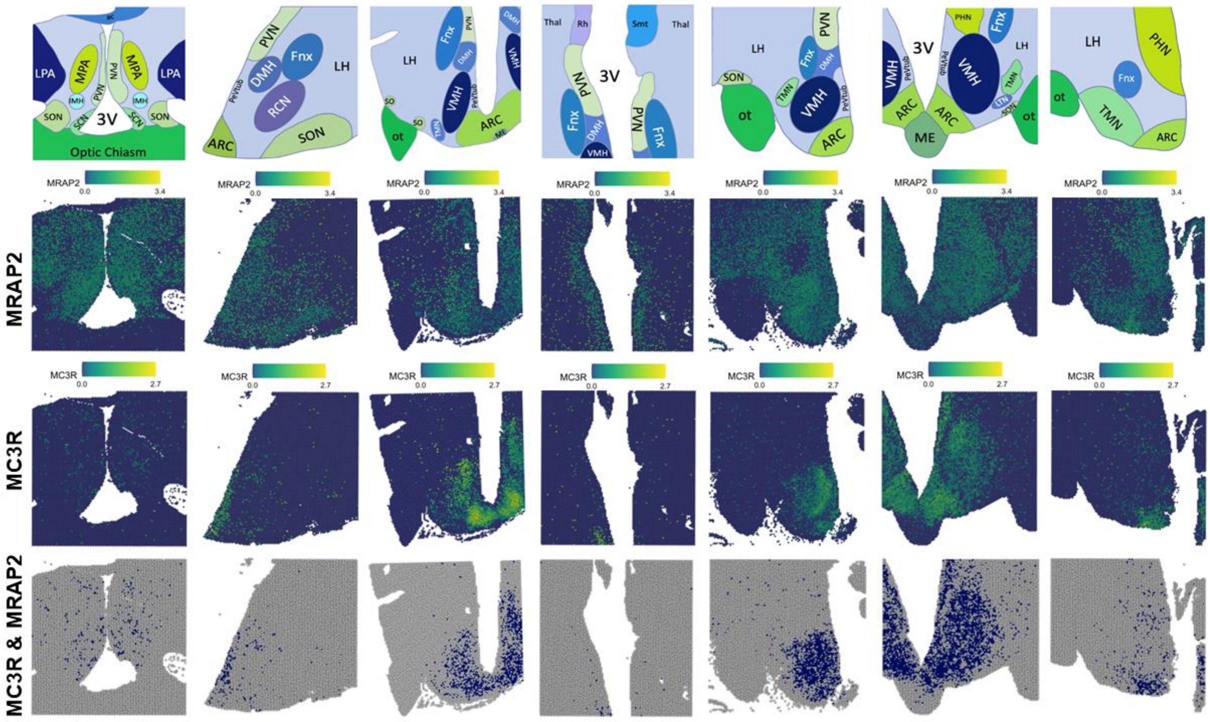


Spatial transcriptomic data showing log-normalized expression of MRAP2 and MC3R in the human hypothalamus. The cartoon shows the major regions of the hypothalamus (ARC, arcuate nucleus; DMH, dorsomedial hypothalamus; Fnx, fornix; LH, lateral hypothalamus, LTN, lateral tuberal nucleus; ME, medial eminence; MPA, medial preoptic area; PeVtub, periventricular nucleus; PHN, posterior hypothalamic nucleus; SCN, suprachiasmatic nucleus; SON, supraoptic nucleus; TMN, tuberomammillary nucleus, VMH, ventromedial nucleus of the hypothalamus), with areas in which MRAP2 and MC3R are co-expressed shown at the bottom (indicated in blue).

**Figure S2 MC3R plasmids express, traffic and signal normally**

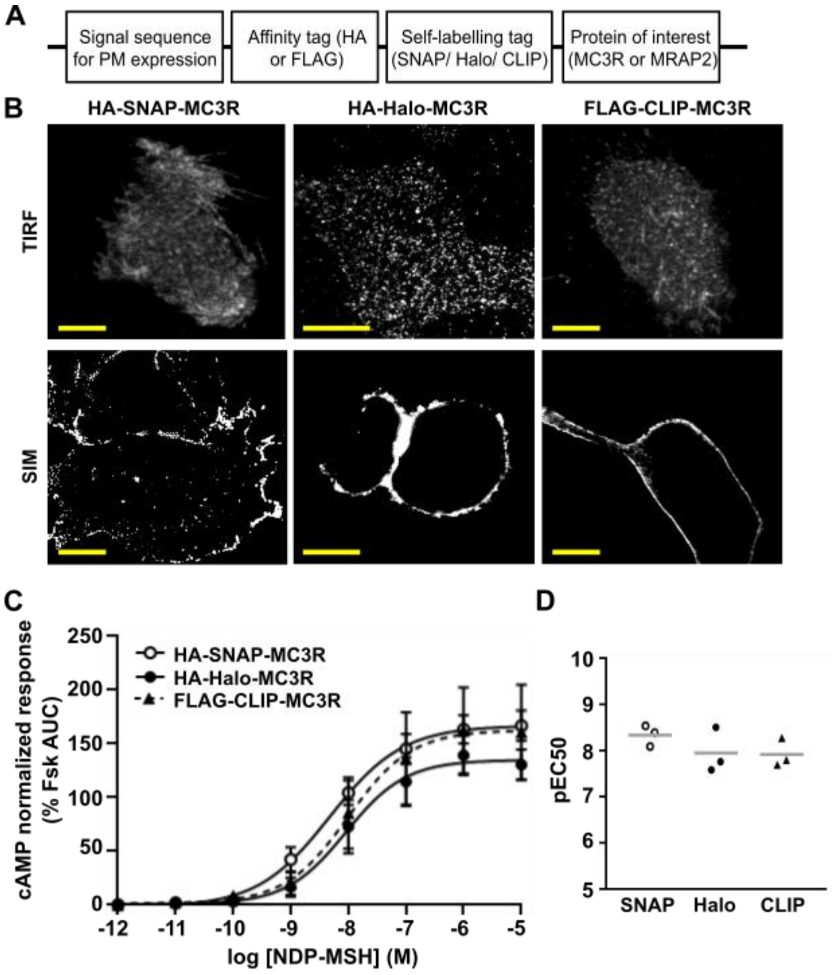


(**A**) Schematic showing the construction of the MC3R and MRAP2 plasmids with a signal sequence, affinity tag (HA or FLAG), self-labeling tag (SNAP, Halo or CLIP) and the protein of interest, (**B**) Imaging showing expression of the three MC3R constructs at the cell surface by TIRF and SIM. Scale, 5 µm. (**C**) MC3R-induced cAMP responses measured by Glosensor in cells transfected with the three MC3R plasmids and (**D**) pEC50. AUC was used to generate a dose-response and expressed relative to basal responses. N=3. Statistical analyses were performed by two-way ANOVA with Sidak's multiple-comparisons test.

**Figure S3 MRAP2 plasmids express and signal normally**

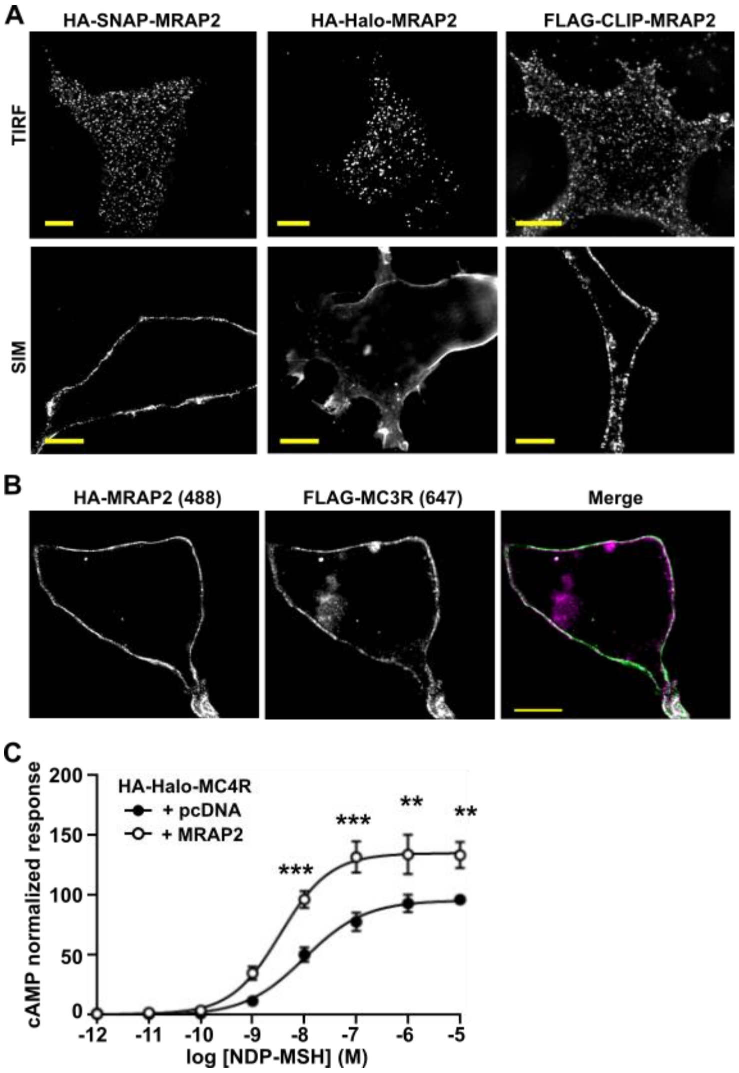


(**A**) Imaging showing expression of the three MRAP2 constructs at the cell surface by TIRF and SIM. Scale, 5 µm. (**B**) SIM imaging showing colocalization of FLAG-CLIP-MC3R and Halo-HA-MRAP2 at the cell surface. Scale, 5 µm. (**C**) MC4R-induced cAMP responses measured by Glosensor in cells transfected with pcDNA or MRAP2, confirming that the MRAP2 plasmids are able to enhance MC4R signaling. AUC was measured and expressed relative to the pcDNA maximal response. N=4. Statistical analyses were performed by two-way ANOVA with Sidak's multiple-comparisons test. ***$p<0.001$, **$p<0.01$.

**Figure S4** **MC3R and MRAP2 interact in a 1:1 stoichiometry**

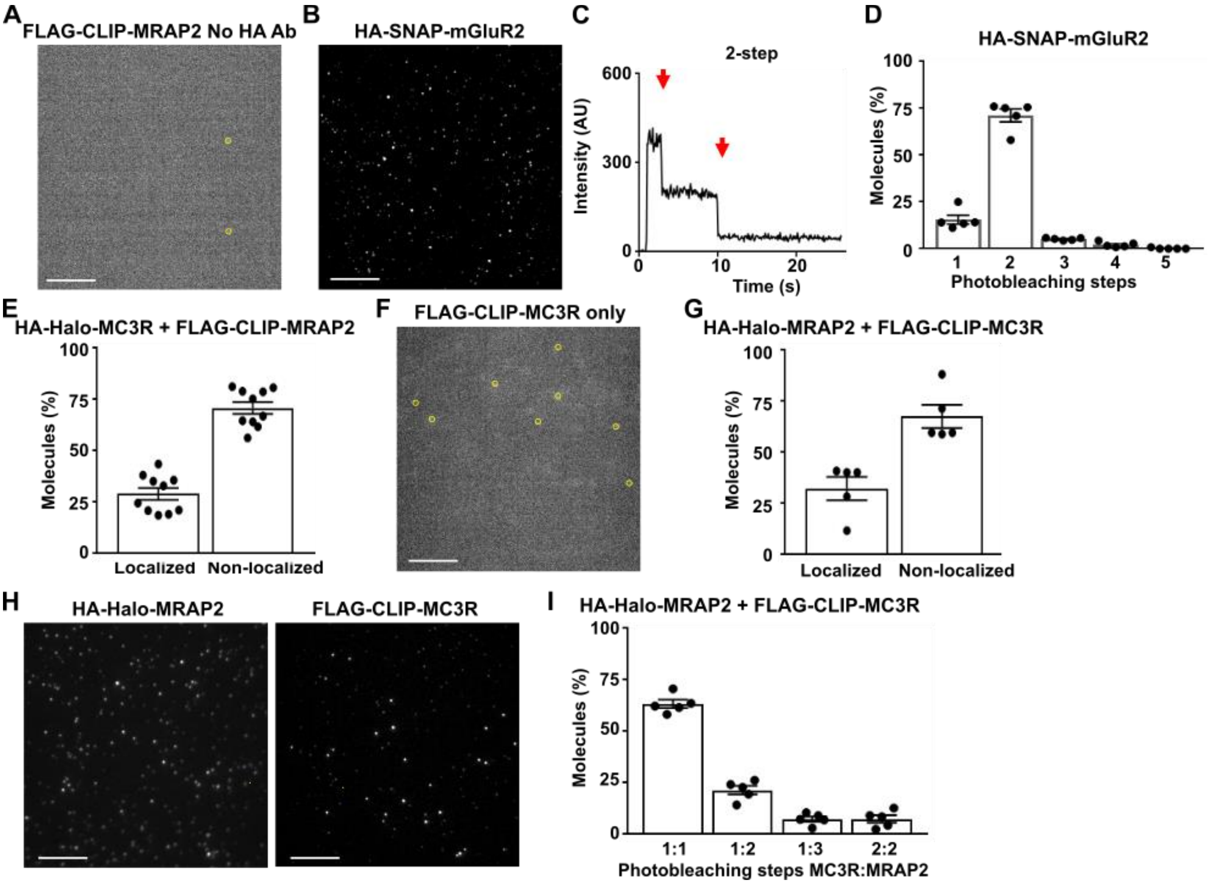


(**A**) Representative image showing little background fluorescence in the absence of anti-HA antibodies. Background fluorescent spots are shown with yellow circles. (**B**) Representative single-molecule fluorescence image of HA-SNAP-mGluR2 with (**C**) examples of single-molecule fluorescence traces with photobleaching steps (red arrows). (**D**) Proportion of molecules with 1 to 5 bleaching steps. N= 1425 molecules from 5 movies. (**E**) Proportion of molecules in two-color SiMPull that are colocalized in cells transfected with HA-Halo-MC3R and FLAG-CLIP-MRAP2. (**F**) Cells transfected with FLAG-CLIP-MC3R only, showing negligible background fluorescence. Background fluorescent spots are shown with yellow circles. (**G**) Proportion of molecules in two-color SiMPull that are colocalized in cells transfected with HA-Halo-MRAP2 and FLAG-CLIP-MC3R. (**H**) Representative two-color SiMPull images of HA-Halo-MRAP2 and FLAG-CLIP-MC3R and (**I**) Photobleaching step analysis from colocalized spots. As in Figure 2, the majority of colocalized spots show 1:1 stoichiometry. N=537 molecules from 5 movies. Scale, 10 μm for all.

**Figure S5 MRAP2 does not interact with SSTR3**

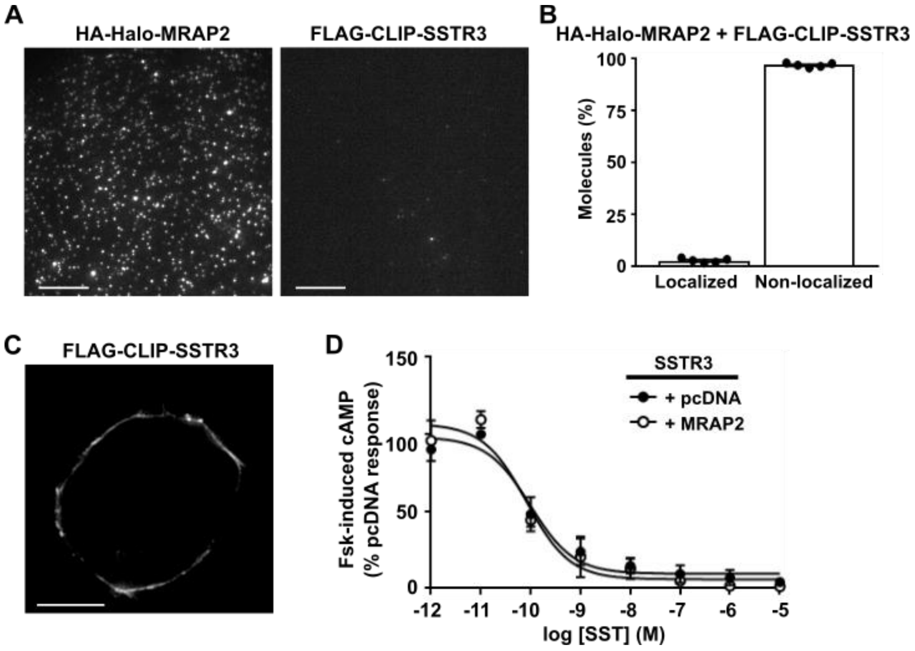


(**A**) Representative two-color SiMPull images of HA-Halo-MRAP2 and FLAG-CLIP-SSTR3, with (**B**) Quantification of the proportion of molecules that are co-localized. N=1790 molecules from 5 movies. Scale, 10 μm. (**C**) SIM image showing FLAG-CLIP-SSTR3 is expressed at the cell surface. Scale, 5 μm. (**D**) FLAG-CLIP-SSTR3-mediated reductions in forskolin (Fsk)-induced cAMP in the presence of pcDNA or MRAP2. N=3.

**Figure S6** **MRAP2 enhances MC3R activity when equal DNA concentrations are transfected**

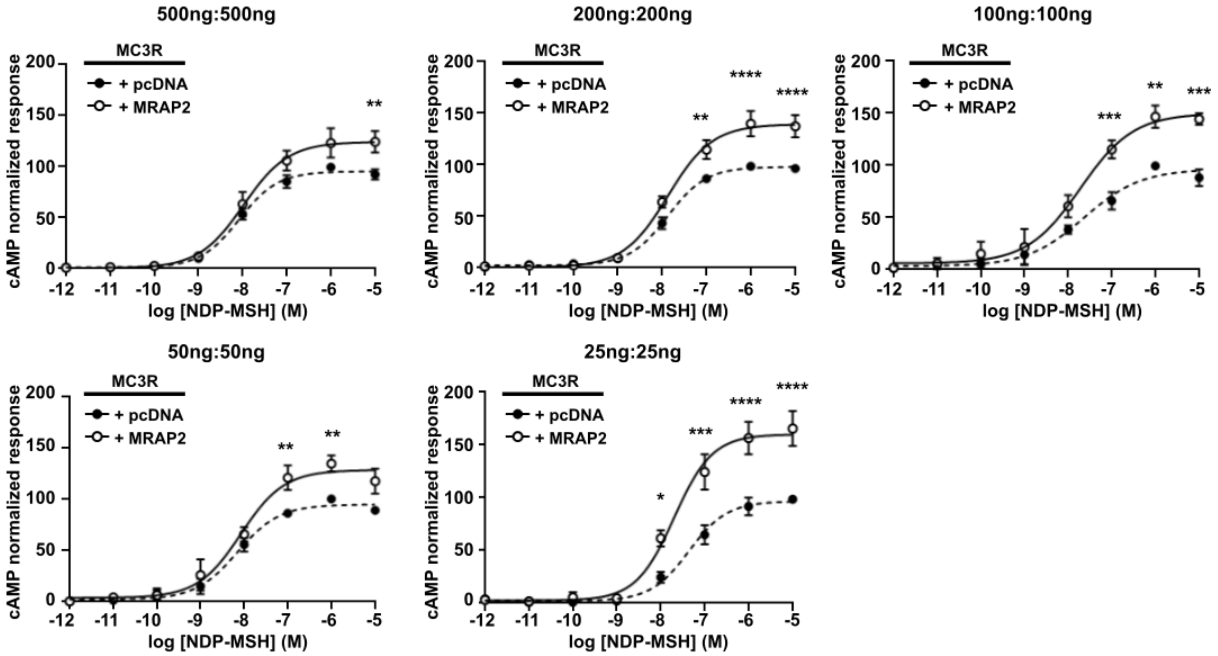


Examination of cAMP responses measured by Glosensor in cells transfected with different total amounts of MC3R with pcDNA or MRAP2. In all assays the same DNA concentration of MC3R and pcDNA or MRAP2 were transfected. AUC was measured and responses expressed relative to the pcDNA maximal response. MRAP2 enhanced MC3R-driven signaling in all assays. N=4. Statistical analyses were performed by two-way ANOVA with Sidak’s test. ****$p<0.0001$, ***$p<0.001$, **$p<0.01$. *$p<0.05$.

**Figure S7 AlphaFold2 models showing predicted interactions between MC3R and MRAP2**

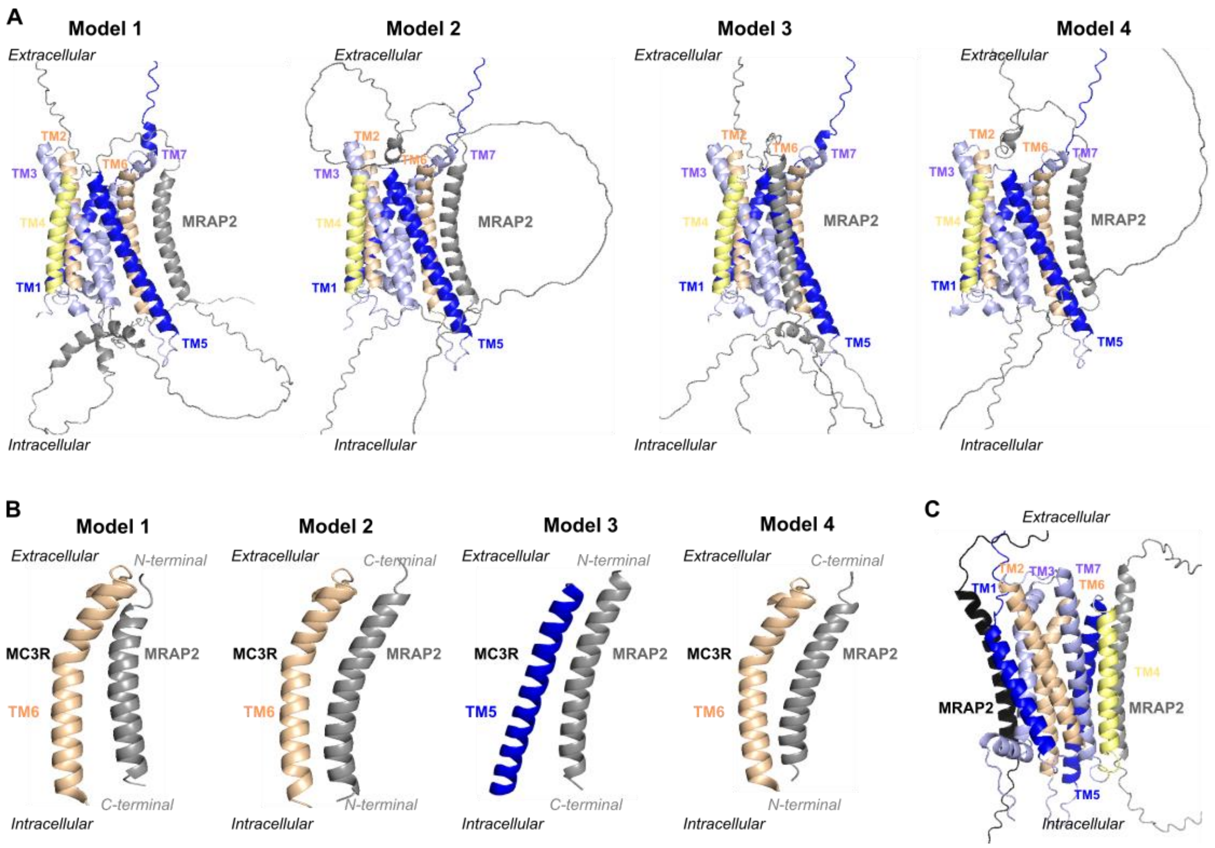


(**A**) Predicted structural models of MC3R and MRAP2 interactions. Model 1 had the highest confidence score, fewer unstructured regions and resembled the MRAP1-MC2R cryo-EM structure (*1*). (**B**) The four models predicted MRAP2 interacts with TM5-TM7, which are known to be important for receptor activation and G protein coupling to MC3R (*2*). Two models predicted MRAP2 inserts into membranes with an extracellular N-terminal region, while two predicted an intracellular N-terminal region. (**C**) Predicted structural model between MC3R and two MRAP2 proteins. All structures similarly predicted the two MRAP2 proteins to bind at distinct sites. No structures predicted MRAP2 dimers.

**Figure S8 Effect of MRAP2 alanine mutations on MRAP2 and MC3R expression**

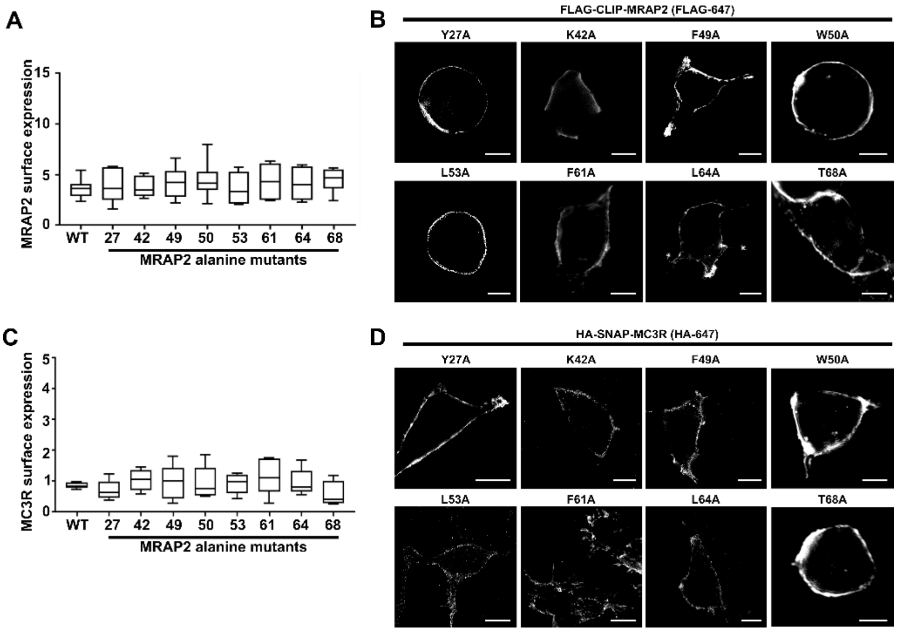


(**A**) Surface expression of MRAP2 assessed in cells transfected with the FLAG-MRAP2 variants. Assays were performed in 96-well plates with primary antibody FLAG and secondary antibody anti-mouse-647. Values were normalized to pcDNA transfected cells. (**B**) MRAP2 surface expression in non-permeabilized cells measured by SIM. Scale, 5 µm. (**C**) Surface expression of MC3R in cells transfected with the HA-HALO-MC3R and FLAG-MRAP2 variants. Assays were performed in 96-well plates with primary antibody HA and secondary antibody anti-mouse-647. Values were normalized to pcDNA transfected cells. (**D**) MC3R surface expression in non-permeabilized cells transfected with the eight alanine variants, measured by SIM. Scale, 5 µm.

**Figure S9 Dyngo-4a blocks MC3R internalization**

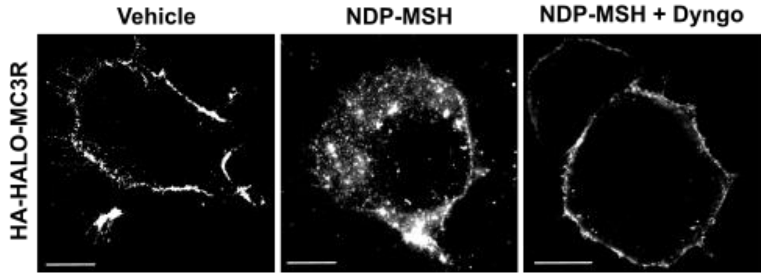


(**A**) Assessment of the effect of Dyngo-4a on cells expressing HA-HALO-MC3R. Cells were exposed to Vehicle, NDP-MSH or agonist with Dyngo-4a and incubated with HA antibody. The HA antibody is only present at the cell surface in vehicle treated cells, while internalization is observed in cells exposed to agonist. In cells pre-treated with Dyngo-4a, there is no internalization observed after 30 minutes exposure to agonist. Scale, 5 μm.

**Figure S10 Effect of MRAP2 human variants on MC3R cell surface expression**

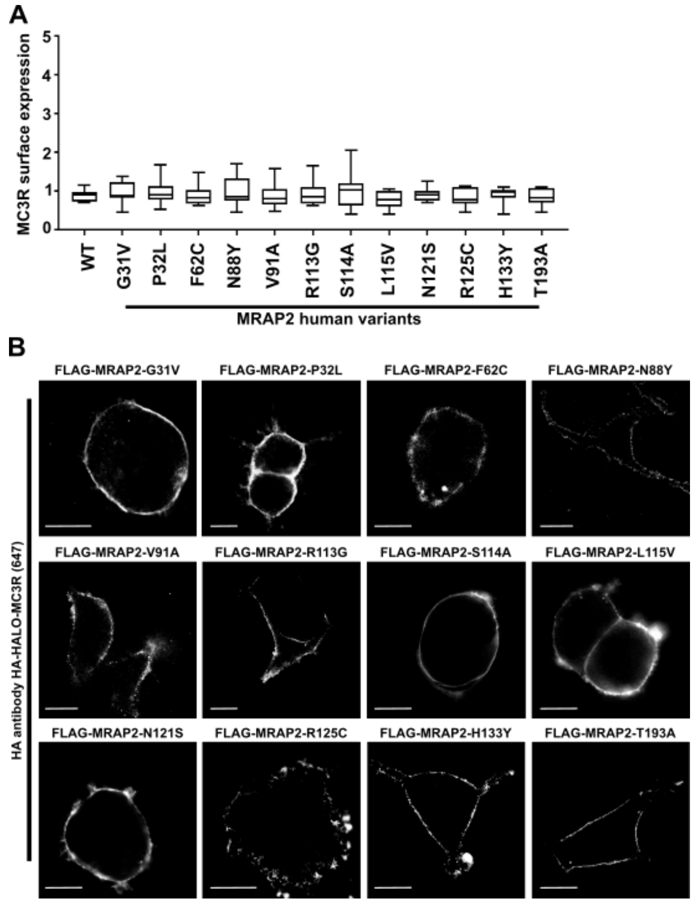


(**A**) Surface expression of MC3R in cells transfected with HA-HALO-MC3R and FLAG-MRAP2 variants. Assays were performed in 96-well plates with primary antibody HA and secondary antibody anti-mouse-647. Values were normalized to pcDNA transfected cells. (**B**) MC3R surface expression in non-permeabilized cells measured by SIM. Scale, 5 μm.

**Table S1 Top 15 MC3R-positive and top 15 MC4R-positive clusters in the human HYPOMAP dataset**

| | | | % cells/ cluster | | | Average log normalized expression/ cluster | | | % co-expressing cells/ cluster | |
|---|---|---|---|---|---|---|---|---|---|---|
| **Cluster name** | **Predicted region** | **Top marker genes** | **MRAP2** | **MC3R** | **MC4R** | **MRAP2** | **MC3R** | **MC4R** | **MC3R & MRAP2** | **MC4R & MRAP2** |
| C4-390 Mid-2 GABA-GLU-3 PGR TAC1 | ARC | KISS1\|NR5A2\|SKOR2\|PGR | 46.94 | 27.76 | 20.82 | 0.41 | 0.17 | 0.12 | 16.73 | 10.61 |
| C4-388 Mid-2 GABA-GLU-3 PGR CALCR | ARC | PGR\|CALCR\|NR5A2\|LHX4 | 37.06 | 23.08 | 4.9 | 0.12 | 0.06 | 0.01 | 12.59 | 2.8 |
| C4-64 Mid-1 GABA-1 NR5A2 | Periventricular | SLC6A3\|NR5A2\|DLK1\|MC3R | 19.23 | 21.15 | 1.92 | 0.21 | 0.31 | 0.03 | 4.81 | 0 |
| C4-391 Mid-2 GABA-GLU-3 PGR TAC3 | ARC | KISS1\|SKOR2\|UGT2B7\|TAC3 | 26.82 | 14.8 | 15.08 | 0.21 | 0.08 | 0.11 | 6.7 | 5.59 |
| C4-161 Mid-1 GABA-6 IL13RA1 GHRH | ARC | GHRH\|GAL\|ADGRF4\|GHSR | 8.56 | 12.83 | 0 | 0.07 | 0.1 | 0 | 0.53 | 0 |
| C4-345 Mid-2 GLU-2 ARHGAP42 COL15A1 | VMH | NR5A1\|COL15A1\|FEZF1\|CCBE1 | 27.8 | 11.5 | 12.99 | 0.23 | 0.09 | 0.13 | 5.32 | 6.07 |
| C4-375 Mid-2 GABA-GLU-3 POMC ANKRD30A | ARC | SOX3\|PGR\|POMC\|TBX3 | 69.75 | 10.92 | 16.25 | 0.65 | 0.04 | 0.09 | 8.96 | 14.01 |
| C4-136 Mid-1 GABA-5 GAL PGR | ARC | GAL\|MBNL3\|NTS\|PGR | 24.3 | 8.41 | 11.53 | 0.16 | 0.06 | 0.08 | 1.87 | 2.49 |
| C4-376 Mid-2 GABA-GLU-3 PDGFD PGR | ARC | PGR\|GABRE\|ALDH1A1\|VGLL3 | 26.05 | 7.78 | 11.08 | 0.24 | 0.02 | 0.06 | 3.89 | 5.99 |
| C4-374 Mid-2 GABA-GLU-3 POMC CALCR | ARC | CALCR\|POMC\|WIF1\|PGR | 33.69 | 7.28 | 2.7 | 0.27 | 0.03 | 0.01 | 4.31 | 1.35 |
| C4-349 Mid-2 GLU-2 SLC22A10 CCBE1 | VMH | SLC22A10\|PGR\|DNAH11\|ADAMTSL1 | 50 | 6.8 | 9.71 | 0.45 | 0.05 | 0.07 | 4.37 | 4.85 |
| C4-385 Mid-2 GABA-GLU-3 PGR DNAH11 | ARC | SKOR2\|KISS1\|DNAH11\|VGLL3 | 22.17 | 6.6 | 17.45 | 0.14 | 0.02 | 0.08 | 2.83 | 5.66 |
| C4-387 Mid-2 GABA-GLU-3 PGR TRPC6 | ARC | PGR\|TBX3\|PGR-AS1\|NR5A2 | 45.37 | 6.32 | 19.19 | 0.28 | 0.03 | 0.11 | 2.71 | 10.38 |
| C4-207 Pre-2 GABA-4 SATB2 TNS3 | MPOA | SATB2\|COL15A1\|EGFLAM\|ZIC2 | 39.51 | 5.85 | 5.37 | 0.38 | 0.03 | 0.05 | 4.88 | 2.44 |
| C4-76 Mid-1 GABA-2 CLMP MBNL3 | SCN | NR2F2-AS1\|ARHGAP36\|SP9\|Z96074.1 | 63.19 | 5.56 | 4.17 | 0.43 | 0.02 | 0.03 | 5.56 | 2.08 |
| C4-303 Mid-3 GLU-3 SLITRK6 FBN2 | MAM | FBN2\|TACR3\|SIM1\|OTP | 52.68 | 0 | 41.07 | 0.28 | 0 | 0.23 | 0 | 25 |
| C4-144 Mid-1 GABA-5 RORB GLI3 | MPOA | GLI3\|TPTE\|GAL\|HMX3 | 58.11 | 0 | 31.08 | 0.49 | 0 | 0.23 | 0 | 21.62 |
| C4-328 Mid-2 GLU-1 SOX14 CYP19A1 | MPOA | SKOR2\|CYP19A1\|FAM9B\|QRFPR | 56.45 | 0 | 27.96 | 0.39 | 0 | 0.18 | 0 | 17.2 |
| C4-194 Pre-2 CHOL-1 BMPR1B | LPOA | SLC5A7\|COL6A5\|CHAT\|LHX8 | 37.7 | 0 | 27.16 | 0.29 | 0 | 0.31 | 0 | 13.42 |
| C4-306 Mid-3 GLU-3 CD36 LMCD1 | MAM | OTP\|KCNH8\|LMCD1\|STK32B | 52.9 | 0 | 26.09 | 0.46 | 0 | 0.2 | 0 | 15.22 |
| C4-206 Pre-2 GABA-4 LHX6 NR0B1 | NA | NPY\|SHISAL2B\|NR0B1\|LHX6 | 66.23 | 0 | 24.68 | 0.57 | 0 | 0.13 | 0 | 20.78 |
| C4-396 Mid-2 GLU-4 WNT7B PRRX1 | NA | LEF1\|WIF1\|RSPO3\|TRABD2B | 18.02 | 0 | 22.52 | 0.12 | 0 | 0.15 | 0 | 5.41 |
| C4-406 Mid-2 GLU-4 EYA4 NPNT | TMN | WIF1\|LEF1\|TBX3\|VEGFC | 33.2 | 3.86 | 21.24 | 0.23 | 0.01 | 0.13 | 2.32 | 10.81 |
| C4-18 Pre-1 GABA-1 SEMA3C CAV1 | NA | CAV1\|PROK2\|CXCL14\|CALCRL | 16.67 | 0 | 20.6 | 0.28 | 0 | 0.27 | 0 | 4.86 |
| C4-171 Pre-2 GABA-1 IL1RAPL2 SST | LPOA | MOXD1\|IL1RAPL2\|LHX6\|NXPH2 | 62.34 | 0 | 20.08 | 0.81 | 0 | 0.21 | 0 | 16.32 |
| C4-29 Pre-1 GABA-1 PRKCH PWWP3B | NA | TAC3\|SLC22A10\|SCML4\|EBF1 | 82.29 | 0 | 20 | 0.77 | 0 | 0.1 | 0 | 17.14 |
| C4-382 Mid-2 GABA-GLU-3 LEF1 IL1RAPL2 | TMN | LEF1\|IL1RAPL2\|CCN3\|ANKRD30A | 47.25 | 0 | 19.72 | 0.37 | 0 | 0.12 | 0 | 11.01 |
| C4-348 Mid-2 GLU-2 SLC22A10 WDR64 | VMH | NR2F2-AS1\|SLC22A10\|SATB1-AS1\|CCBE1 | 44.53 | 2.73 | 19.53 | 0.43 | 0.03 | 0.13 | 0.78 | 12.11 |
| C4-118 Mid-1 GABA-4 NTS ABCG2 | MPOA | TMEM114\|ESR1\|FAM9B\|PGR | 78.88 | 0 | 19.4 | 0.7 | 0 | 0.08 | 0 | 18.53 |

ARC, arcuate nucleus; MPOA, medial preoptic area; NA, nucleus accumbens; SCN, suprachiasmatic nucleus; TMN, Tuberomammillary nucleus; VMH, ventromedial nucleus of the hypothalamus.

**Table S2 Expression plasmids used in this manuscript**

| Plasmid name | | Information | | Source |
|---|---|---|---|---|
| cAMP Glosensor-22F | | cAMP sensor | | Promega |
| ss-HA-Halo-MC3R | | N-terminal signal peptide from mGluR5, followed by HA, HALO and human MC3R | | This manuscript |
| ss-HA-SNAP-MC3R | | N-terminal signal peptide from mGluR5, followed by HA, SNAP and human MC3R | | This manuscript |
| ss-FLAG-CLIP-MC3R | | N-terminal signal peptide from mGluR5, followed by HA, CLIP and human MC3R | | This manuscript |
| ss-HA-Halo-MRAP2 | | N-terminal signal peptide from mGluR5, followed by HA, Halo and human MC3R | | This manuscript |
| ss-HA-SNAP-MRAP2 | | N-terminal signal peptide from mGluR5, followed by HA, SNAP and human MC3R | | This manuscript |
| ss-FLAG-CLIP-MRAP2 | | N-terminal signal peptide from mGluR5, followed by HA, CLIP and human MC3R | | This manuscript |
| ss-HA-Halo-mGluR2 | | Used as template for ss-HA-Halo-MC3R | | Joshua Levitz, Weill Cornell Medicine |
| ss-HA-SNAP-mGluR2 | | Used as template for ss-HA-SNAP-MC3R | | Joshua Levitz, Weill Cornell Medicine |
| ss-FLAG-CLIP-mGluR2 | | Used as template for ss-FLAG-CLIP-MC3R | | Joshua Levitz, Weill Cornell Medicine |
| ss-HA-Halo-MC4R | | N-terminal signal peptide from mGluR5, followed by HA, HALO and human MC3R | | This manuscript |
| ss-FLAG-CLIP-SSTR3 | | N-terminal signal peptide from mGluR5, followed by HA, HALO and human SSTR3 | | This manuscript |
| SSTR3-Tango | | Used as a template for SSTR3 constructs | | Bryan Roth, Addgene plasmid #66504 |
| MC3R-Tango | | Used as a template for MC3R constructs | | Bryan Roth, Addgene plasmid #66429 |
| MRAP2-3xFLAG | | Used as a template for MRAP2 constructs and in most assays | | Julien Sebag, University of Iowa |
| B-arrestin2-mYFP | | SIM | | Robert Lefkowitz, Addgene plasmid #36917 |
| Nluc-Arr2 | | BRET | | Steve Hill, University of Nottingham |
| Venus-Kras | | BRET | | Nevin Lambert, Augusta University |
| Rab5-Venus | | SIM | | Nevin Lambert, Augusta University |
| LgC-MC3R | | NanoBiT | | This manuscript |
| SmC-MC3R | | NanoBiT | | This manuscript |
| LgC-MRAP2 | | NanoBiT | | This manuscript |
| SmC-MRAP2 | | NanoBiT | | This manuscript |
| MRAP2-3xFLAG-Y27A | | Glosensor, SIM, cell surface expression | | This manuscript |
| MRAP2-3xFLAG-K42A | | Glosensor, SIM, cell surface expression | | This manuscript |
| MRAP2-3xFLAG-F49A | | Glosensor, SIM, cell surface expression | | This manuscript |
| MRAP2-3xFLAG-W50A | | Glosensor, SIM, cell surface expression | | This manuscript |
| MRAP2-3xFLAG-L53A | | Glosensor, SIM, cell surface expression | | This manuscript |
| MRAP2-3xFLAG-F61A | | Glosensor, SIM, cell surface expression | | This manuscript |
| MRAP2-3xFLAG-L64A | | Glosensor, SIM, cell surface expression | | This manuscript |

| | | | | |
|---|---|---|---|---|
| MRAP2-3xFLAG-T68A | | Glosensor, SIM, cell surface expression | | This manuscript |
| ss-HA-Halo-MC3R-T245A | | Glosensor, SIM, cell surface expression | | This manuscript |
| ss-HA-Halo-MC3R-L260A | | Glosensor, SIM, cell surface expression | | This manuscript |
| ss-HA-Halo-MC3R-P272A | | Glosensor, SIM, cell surface expression | | This manuscript |
| MRAP2-3xFLAG-G31V | | Glosensor, SIM, cell surface expression | | This manuscript |
| MRAP2-3xFLAG-P32L | | Glosensor, SIM, cell surface expression | | This manuscript |
| MRAP2-3xFLAG-F62C | | Glosensor, SIM, cell surface expression | | This manuscript |
| MRAP2-3xFLAG-N88Y | | Glosensor, SIM, cell surface expression | | This manuscript |
| MRAP2-3xFLAG-V91A | | Glosensor, SIM, cell surface expression | | This manuscript |
| MRAP2-3xFLAG-R113G | | Glosensor, SIM, cell surface expression | | This manuscript |
| MRAP2-3xFLAG-S114A | | Glosensor, SIM, cell surface expression | | This manuscript |
| MRAP2-3xFLAG-L115V | | Glosensor, SIM, cell surface expression | | This manuscript |
| MRAP2-3xFLAG-N121S | | Glosensor, SIM, cell surface expression | | This manuscript |
| MRAP2-3xFLAG-R125C | | Glosensor, SIM, cell surface expression | | This manuscript |
| MRAP2-3xFLAG-H133Y | | Glosensor, SIM, cell surface expression | | This manuscript |
| MRAP2-3xFLAG-T193A | | Glosensor, SIM, cell surface expression | | This manuscript |

**Table S3 Identification of possible contacts between MRAP2 and MC3R in AlphaFold2 models**

| | | MC3R | | | | |
|---|---|---|---|---|---|---|
| **MRAP2** | **Region[a]** | **Rank 1** | **Rank 2** | **Rank 3** | **Rank 4** | **No. of models[b]** |
| **W23** | EC/IC | | N68 | | | 1 |
| **E26** | EC/IC | D117 | | | | 1 |
| **Y27** | EC/IC | F281 | K239, G240 | D121 | Y299, R302 | **4** |
| **I30** | EC/IC | | Q233, H235 | | S236 | 2 |
| **K39** | EC/IC | F34 | | | | 1 |
| **A40** | EC/IC | Y273 | M238 | | | 2 |
| **K42[c]** | TM | P272, Y273 | | M187 | | 2 |
| **Y43** | TM | | V242 | | M238 | 2 |
| **S44** | TM | | F211 | | | 1 |
| **I45** | TM | | | M187 | | 1 |
| **V46** | TM | | | V190 | T245 | 2 |
| **F49** | TM | L260 | | C191, M195 | G249 | **3** |
| **W50[c]** | TM | P257 | T245, L248, G249 | | L248 | **3** |
| **L53[c]** | TM | A256 | | A198 | I252 | **3** |
| **F61[c]** | TM | | V263 | T205 | F197 | **3** |
| **L64[c]** | TM | T245 | L260 | L206 | L260 | **4** |
| **L67** | TM | F211, R215 | | | | 1 |
| **T68** | TM | | Y267 | H209 | | **2** |
| **F90** | IC/EC | | D117 | | | 1 |
| **R113** | IC/EC | P225 | | | | 1 |
| **I120** | IC/EC | | | Q223 | | 1 |
| **F169** | IC/EC | S236 | | | | 1 |

The Rank 1 model has fewer unstructured regions and is most similar to the published MRAP1-MC2R structure (*1*). [a]Region refers to the structural location as either: EC, extracellular region, TM, transmembrane helix, IC, intracellular region. Some residues are designated as IC/EC as the orientation of MRAP2 differs in some models (Figure S7). [b]No. of models refers to the total number (out of 4) of structural models that identified a link between the MRAP2 residue and MC3R. MRAP2 residues highlighted in red were mutated to alanine and functionally characterized in this study, while MC3R residues similarly investigated are highlighted in blue. [c]Alanine substitutions that impair MRAP2 enhancement of MC3R signaling (Figure 5).

**Table S4 Densitometry of FLAG-MRAP2 protein with alanine variants**

| Human MRAP2 (relative to WT) | Mean±SEM (N) |
|---|---|
| Y27A | 1.13 ± 0.03 (5) |
| K42A | 1.04 ± 0.06 (5) |
| F49A | 0.97 ± 0.06 (5) |
| W50A | 1.11 ± 0.03 (5) |
| L53A | 0.93 ± 0.07 (5) |
| F61A | 1.04 ± 0.09 (5) |
| L64A | 0.93 ± 0.09 (5) |
| T68A | 1.68 ± 0.19 (5)**** |
| pcDNA | 0.01 ± 0.01 (5)**** |

Densitometry of FLAG-MRAP2 protein from five western blots. MRAP2 variants were investigated in separate batches and protein expression was normalized to the MRAP2-WT control in each experiment. Asterisks indicate significant difference to WT. ****p<0.0001 compared to MRAP2-WT.

**Table S5 Identification of possible contacts between MC3R and MRAP2 residues mutated in overweight and/or obese individuals**

| Variant | Rank 1 | Rank 2 | Rank 3 | Rank 4 |
|---|---|---|---|---|
| **G31V** | E29 | None | V33 | None |
| **P32L** | S34 | None | S34 | None |
| **F62C** | None | I58, L66 | None | None |
| **N88Y** | None<br>Mutant forms new contact with S89. | R84, S92. Mutant forms new contacts with F85, R86. | None | S92<br>Mutant forms new contact with R84. |
| **V91A** | None | None | None | F94 |
| **R113G** | Link to MC3R P225.<br>Mutant loses contact. | None | E111 | None |
| **S114A** | Link to MC3R N159.<br>Mutant loses contact. | None | None | None |
| **L115V** | None | None | None | None |
| **N121S** | H117, C118, R125, N171 | None | H117, C118, R125, E124.<br>Mutant loses H117 contact. | None |
| **R125C** | N121, A129 | None | N121, E122 | None |
| **H133Y** | K130 | None | K130<br>Mutant loses contact. | None |
| **T193A** | None | None | None | None |

Contacts are in MRAP2 (i.e. intramolecular) unless otherwise stated. The predicted consequences of MRAP2 variants are highlighted in red. Those in which there is no red text predicted no structural effect of the MRAP2 variant.
